# The medial entorhinal cortex encodes multisensory spatial information

**DOI:** 10.1101/2024.01.09.574924

**Authors:** Duc Nguyen, Garret Wang, Yi Gu

## Abstract

Animals employ spatial information in multisensory modalities to navigate their natural environments. However, it is unclear whether the brain encodes such information in separate cognitive maps or integrates all into a single, universal map. We addressed this question in the microcircuit of the medial entorhinal cortex (MEC), a cognitive map of space. Using cellular-resolution calcium imaging, we examined the MEC of mice navigating virtual reality tracks, where visual and auditory cues provided comparable spatial information. We uncovered two cell types: “unimodality cells” and “multimodality cells”. The unimodality cells specifically represent either auditory or visual spatial information. They are anatomically intermingled and maintain sensory preferences across multiple tracks and behavioral states. The multimodality cells respond to both sensory modalities with their responses shaped differentially by auditory and visual information. Thus, the MEC enables accurate spatial encoding during multisensory navigation by computing spatial information in different sensory modalities and generating distinct maps.

## Introduction

We have a fairly good understanding on how spatial navigation utilizes visual information. However, navigation often incorporates spatial information in multisensory modalities^1,2^, such as navigations of nocturnal animals at nights and visually impaired people on a street. Does the brain encode spatial information in different sensory modalities in separate maps or integrate all into a universal cognitive map? This fundamental question is key for understanding how the brain achieves precise spatial encoding during multisensory navigation. Here we pursued this question by investigating the encoding of visual and auditory spatial information in mouse medial entorhinal cortex (MEC), which is hypothesized to form a “cognitive map” of space^3,4^.

Nocturnal rodents, such as mice and rats, have good night-vision to detect motion and contrast^5^. In laboratory settings, they can perform navigation tasks in environments with complex visual cues^6–8^. Consistently, rodent MEC contains multiple cell types to encode spatial information during navigation in visual-cue-enriched environments^9–11^. On the other hand, rodents also have strong auditory capabilities to distinguish sounds at various frequencies and amplitudes^12–14^. Auditory spatial cues can guide them to successfully navigate in real^15,16^ and virtual environments^17,18^. However, while the MEC responds to non-spatial acoustic cues, such as tones at various frequencies^19,20^, auditory beats^21^, and noise^22^, and exhibits c-fos expression upon auditory stimuli^23^, it is unclear whether the MEC also represents auditory cues that convey spatial information during navigation. If it does, the next important question would be how auditory encoding in the MEC converges with visual encoding of space, as mentioned above.

Some evidence suggests that the MEC could use a universal map for auditory and visual spatial information. MEC responses were identified not only in real physical space, but also in virtual reality environments^24,25^ and pure cognitive spaces constructed by visual scenes/cues^26–28^, sound frequency^20^, odor^29^, and conceptual knowledge^30–32^. These observations suggest that the MEC could extract metric information of physical and abstract spaces regardless of sensory features of metric variables^3,33–36^. Therefore, the MEC could construct a universal map for auditory and visual spatial cues if they both represent valid metric information. For example, in unisensory environments where auditory and visual spatial cues provide metrically-equivalent spatial information, the same set of MEC cells could exhibit similar responses so that only metric but not sensory information is encoded (Figure 1A, a). If both auditory and visual cues provide metric information of a multisensory environment, MEC cell population could unbiasedly represent all cues, regardless of their sensory modalities (Figure 1B, a).

**Figure 1:**
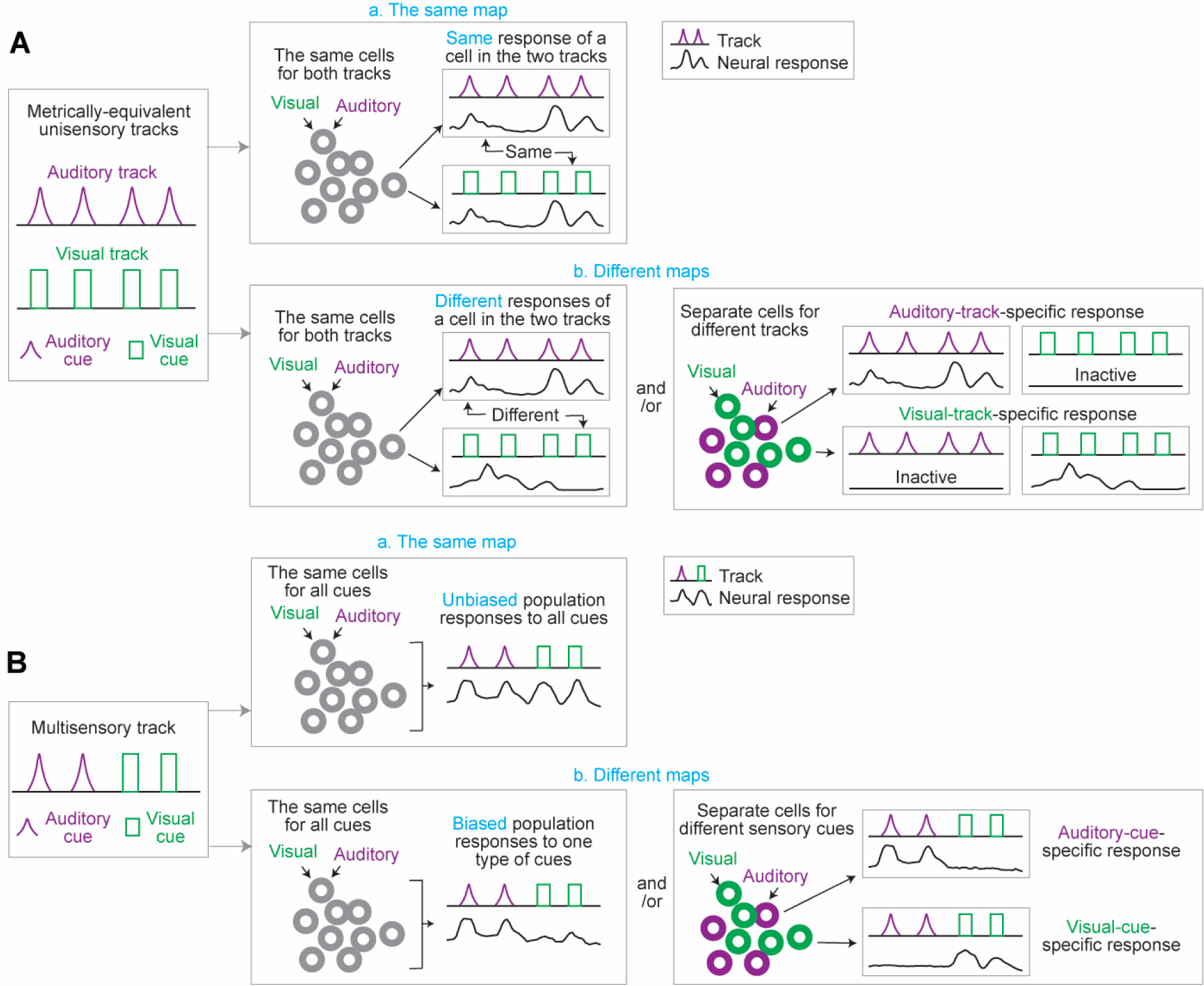
Hypothetical MEC responses in metrically-equivalent tracks and a multisensory track A. MEC response in unisensory auditory and visual tracks with metrically-equivalent spatial information. a: The MEC could use the same map for auditory and visual tracks. The same group of cells show similar responses in the two tracks. b: The MEC could use different maps for auditory and visual tracks. The map could consist of the same group of cells with different responses in the two tracks (left), and/or separate cell groups specifically active in auditory or visual track (right). B. MEC response in a multisensory track with nonoverlapping visual and auditory cues. a: The MEC could use the same map for auditory and visual cues. All cues could be unbiasedly encoded by MEC cell populations. b: The MEC could use different maps for auditory and visual cues. At the population level, there could be biased encoding of one type of sensory cues (auditory cues in this figure, left). There could also be separate cell groups for each type of sensory cues (right).

Conversely, the MEC could use different maps for auditory and visual cues, since previous studies on the hippocampus suggest that sensory feature is an important component of the cognitive map^37^. The hippocampus is reciprocally connected with the MEC^11^ and is also abundant with spatially tuned cells^4^. In spatial environments involving sensory cues, such as sounds and odors, hippocampal neural responses were modulated by both metric location and sensory information^37–40^. It is possible that the MEC cognitive map also encodes sensory information as the hippocampus does. In addition, different patterns of auditory and visual inputs to the MEC could shape different maps. Auditory inputs indirectly arrive at the MEC via the parahippocampal cortex (PHC), retrosplenial cortex (RSC)^41–47^, and medial septum^48^, whereas visual inputs are delivered both indirectly through the PHC/RSC-MEC pathway^42,49,50^ and directly from the visual cortex^45,51,52^. If the MEC indeed uses different maps, in the metrically-equivalent unisensory environments above, the same group of cells could show different responses, and/or separate cells could be recruited by the two environments (Figure 1A, b). Likewise, in the multisensory environment, MEC cell population could exhibit biased responses to a certain type of sensory cues, and there could also be distinct cells specifically encoding each type of cues (Figure 1B, b).

To investigate the above possibilities, we conducted cellular-resolution two-photon imaging to measure MEC calcium dynamics while mice navigated in virtual reality environments with flexibly controlled auditory and visual cues. We first characterized the preferential encoding of the most task-relevant auditory cues in the MEC during navigation in auditory virtual reality (AVR). We further combined AVR with visual virtual reality (VVR) to compare the MEC maps under the above two settings: metrically-equivalent unisensory tracks and a multisensory track (Figure 1). While both auditory and visual cues could guide comparably successful navigation, we consistently demonstrated that these different sensory cues were encoded by different MEC maps at the microcircuit level, suggesting that MEC cognitive map represents not only metric, but also sensory information of space. These results uncover the fundamental neural mechanism of the MEC for multisensory navigation.

## Results

### The MEC exhibits spatial response during auditory-cue-guided navigation

We first investigated MEC neural response during auditory-cue-guided navigation in AVR (Figure 2A), which was operated in darkness and consisted of eight speakers surrounding a head-fixed mouse running on a spherical treadmill^17^. Auditory cues at different frequencies were delivered at fixed locations on a linear track, and water-restricted mice were trained to stop within a hidden reward zone for at least one second to receive a water reward (AVR1, Figure 2B). Spatial distance and orientation of a cue to the mouse were simulated by varying sound intensity and source speakers for the cue at different track locations, respectively (Figures 2B and S1A). GP5.3 mice with stable expression of calcium indicator GCaMP6f in excitatory neurons in MEC layer 2^53,54^ navigated in AVR1, enabling cellular resolution two-photon imaging of MEC calcium dynamics during navigation (Figure 2C). To control for the specificity of behavior and neural response to auditory cues, we conducted a comparable study when mice traversed a no-auditory-cue (NA) track, which was identical to AVR1 but without auditory cues (Figure 2D).

**Figure 2:**
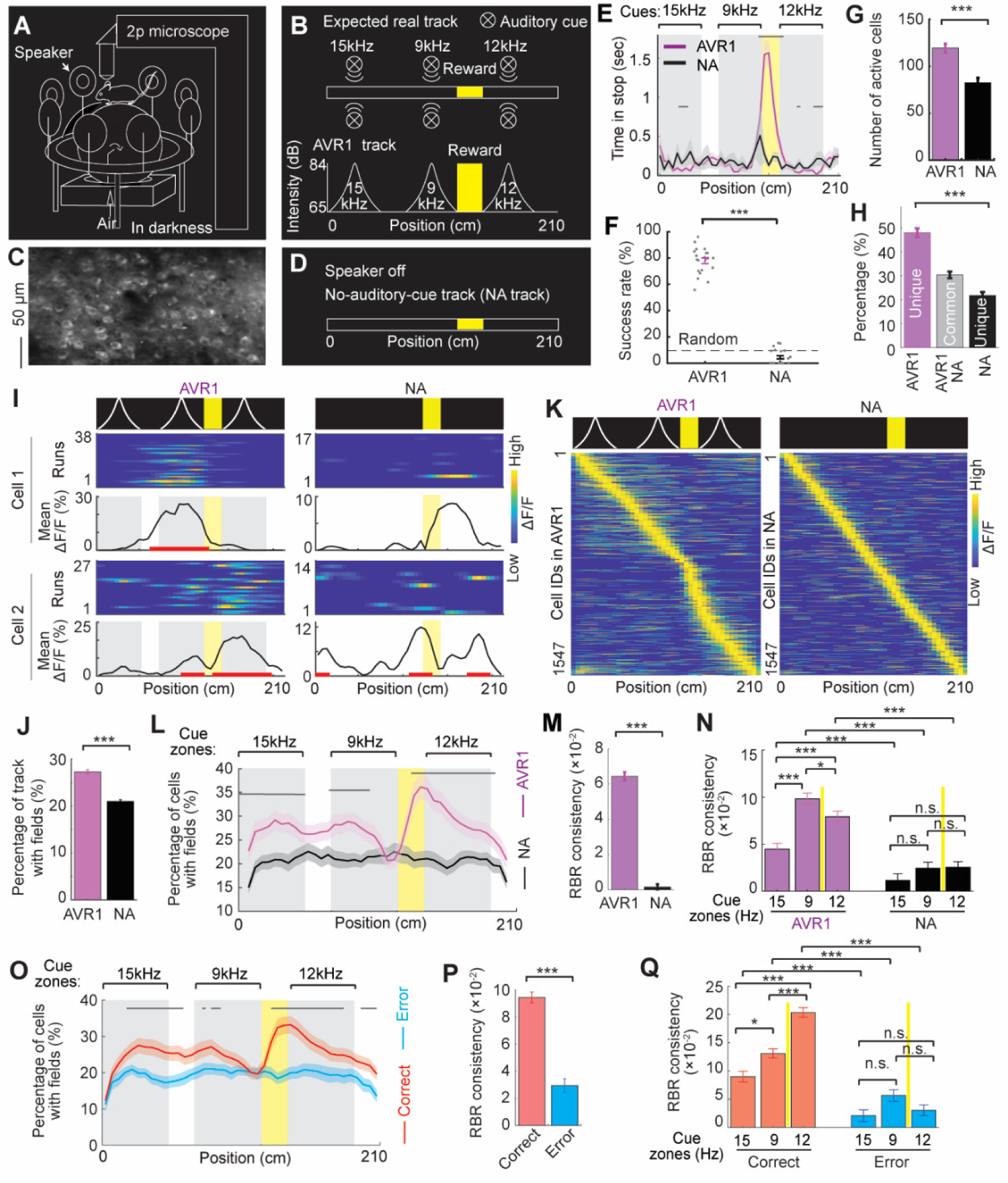
MEC response in AVR A. AVR setup, which was combined with a two-photon (2p) microscope. B. Top: expected real auditory track with audio cues (15, 9, and 12 kHz) delivered on both sides and a reward zone. Bottom: sound intensity of auditory cues changed with AVR1 track locations to simulate the distance effect between auditory cues and the mouse. C. An example imaging field of view. D. A diagram of a no-auditory-cue track (NA). E. Averaged time the mice spent stopping across runs in AVR1 and NA. Grey horizontal lines indicate the areas with significantly different stopping time between AVR1 and NA. The yellow zone indicates reward. Same in following figures. F. Success rate in receiving reward in AVR1 and NA. p = 1.1×10^−19^. The dashed line indicates a random level. G. Number of active cells of the same FOVs in AVR1 and NA. p = 1.4×10^−9^. H. From left to right: the percentage of unique cells in AVR1, common cells in AVR1 and NA, and unique cells in NA. p values for unique cells in AVR1 and in NA: 7.7×10^−11^. I. Two examples of the same cell in AVR1 (left) and in NA (right). For each cell: top: environment template. Middle: cell activity on a run-by-run basis. Bottom: Average activity across runs as a function of track positions (mean ΔF/F) and spatial fields (red lines). Gray and yellow blocks indicate auditory cue areas and reward zone, respectively. Same in following figures. J. Percentage of track with spatial fields of common cells. p= 6.4×10^−27^. K. Activity matrices of common cells in AVR1 and NA. Each row represents one cell, the mean ΔF/F of which within fields were kept as original values and those outside of fields were zero. This mean ΔF/F was further normalized between 0 and 1 and sorted based on its peak location. Same in following figures. L. Percentage of common cells with spatial fields at each spatial bin along AVR1 or NA track. Grey horizontal lines indicate track areas with significantly different percentages between AVR1 and NA. Same in following figures. M. RBR activity consistency of common cells in AVR1 and NA. p = 6.9×10^−98^. N. RBR activity consistency of common cells in individual cue zones in AVR1 and NA. p values for AVR1 and NA in 15 kHz cue: 3.4×10^−4^; 9 kHz cue: 6.1×10^−17^, and 12 kHz cue: 9.0×10^−10^. p values between cues within AVR1: 15 kHz versus 9 kHz: 5.7×10^−10^; 15 kHz versus 12 kHz: 4.1×10^−5^, and 9 kHz versus 12 kHz: 0.0232. Yellow lines indicate the reward zone between cues. Same in following figures. O. Percentage of common cells with spatial fields on AVR1 track in correct and error runs. P. RBR activity consistency of common cells in correct and error runs in AVR1. p = 1.4×10^−20^. Q. RBR activity consistency of common cells in individual cue zones in correct and error runs in AVR1. p values for correct and error runs at 15 kHz cue: 3.6×10^−4^; 9 kHz cue: 4.5×10^−6^, and 12 kHz cue: 3.6×10^−25^. p values between cues within correct runs: 15 kHz versus 9 kHz: 0.0032; 15 kHz versus 12 kHz: 5.4×10^−15^, and 9 kHz versus 12 kHz: 2.5×10^−8^. *p ≤ 0.05, **p ≤ 0.01, ***p ≤0.001, n.s. p > 0.05. Error bars represent mean ± SEM. Data was collected from 5 mice.

In AVR1 but not NA track, mice specifically stopped in the reward zone to receive reward at a high success rate (Figures 2E and 2F). Additionally, correct runs (in which the reward was successfully received) in AVR1 and NA did not lead to higher reward success rates in following runs compared to error runs (in which no reward was received) (Figure S1B), excluding the possibility that the mice identified the reward zone based on other information, such as the number of steps or cues, or elapsed time, rather than auditory cue features.

Different calcium responses of cells in the same set of imaging fields of view (FOVs) in AVR1 and NA support the specificity of the MEC map for auditory spatial information. More cells were active in AVR1 than in NA (Figure 2G). While some cells were commonly active (common cells) in both tracks, the percentage of cells uniquely active (unique cells) in AVR1 was higher than that in NA (Figure 2H). We first focused on common cells and examined their spatial fields, which were defined as track areas consisting of adjacent spatial bins with significantly higher calcium response than random chance (Figure 2I). Spatial fields in the cells showed higher prevalence (the percentage of track covered by fields) in AVR1 than in NA (Figures 2I and 2J). The fields were enriched around three cues in AVR1, but evenly distributed in NA (Figures 2K and 2L). We further investigated spatial activity consistency of the cells on a run-by-run basis (RBR consistency) by calculating the activity correlation between individual runs within the same session. RBR consistency was higher in AVR1 than in NA (Figure 2M). The consistency difference existed at individual cues and higher consistency was observed at the two cues near the reward in AVR1 (Figure 2N). Moreover, the high field abundance around cues and high RBR activity consistency especially at the cues near the reward only occurred in the correct but not the error runs in AVR1 (Figures 2O-Q). Similarly, AVR1 unique cells exhibited specific activity to auditory cues compared to NA unique cells (Figures S1C-E), especially in the correct runs (Figures S1F-H).

Therefore, in the correct but not the error runs, the MEC specifically responded to auditory cues, especially to the most task-relevant ones near the reward. This indicates that during auditory-cue-guided navigation, MEC activity primarily reflects spatial cognition states, rather than sensory perception to auditory cues.

### Metrically-equivalent auditory and visual tracks recruited different MEC cells

We next asked how the encoding of auditory spatial information compares with that of visual information in unisensory auditory and visual tracks with equivalent metric information (Figure 1A). Two pairs of metrically-equivalent tracks, AVR1 versus VVR1, and AVR2 versus VVR2, were created in AVR and VVR (Figures 3A and 3B). Each pair contained tracks with auditory or visual cues at identical locations relative to the hidden reward zone (Figure 3C). AVR2 included all AVR1 cues and an additional cue, so they belonged to the same sensory modality (same-sensory tracks) but delivered different metric information (Figures S2A and S2B). A similar design was followed in VVR2 and VVR1 (Figures S2C and S2D). Hypothetically, similar MEC responses in metrically-equivalent tracks would indicate a universal map for metric information regardless of sensory modality, whereas different responses on metrically-equivalent tracks but similar responses on same-sensory tracks would suggest separate maps for auditory and visual spatial information (Figure 1A).

**Figure 3:**
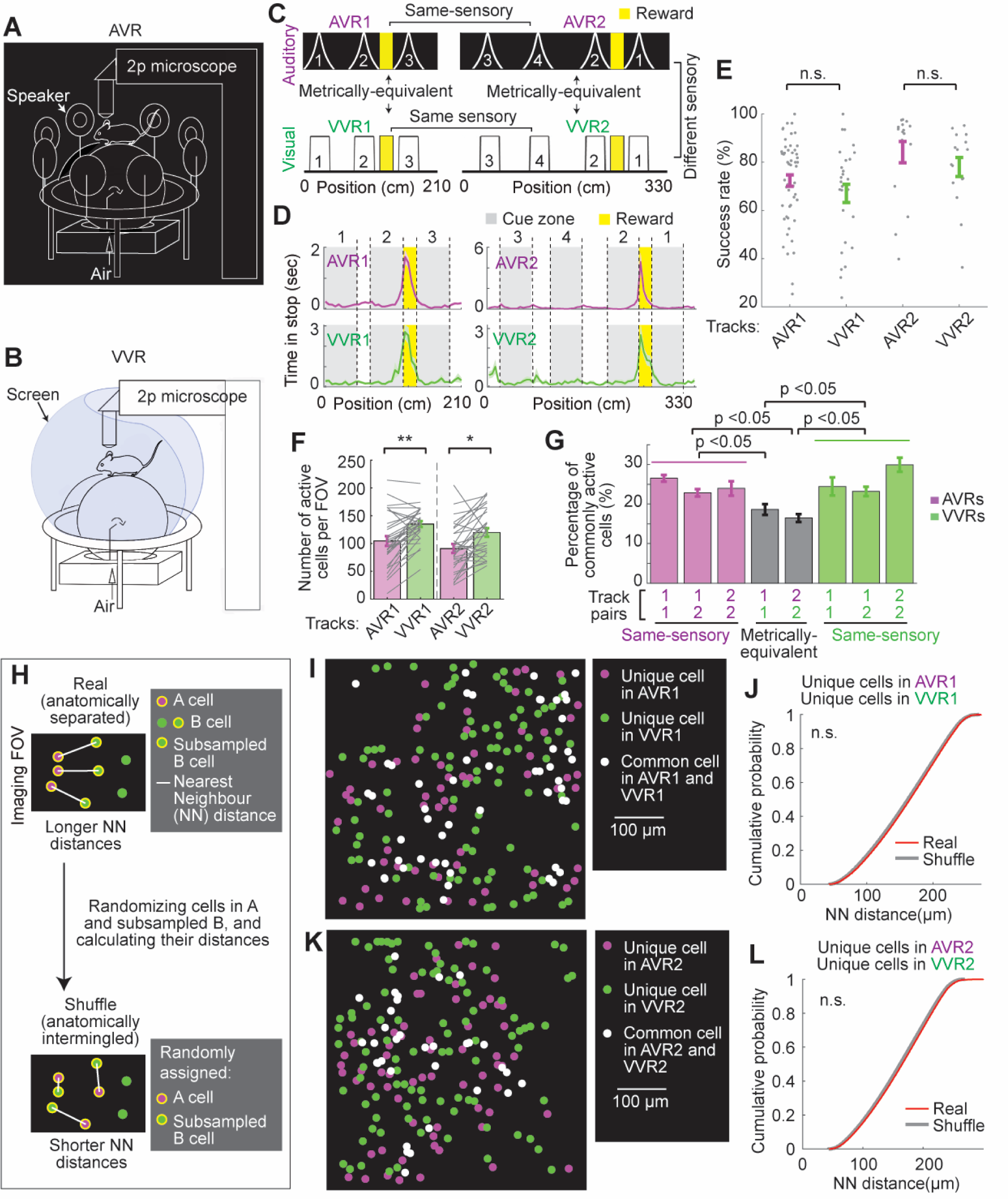
MEC cells in metrically-equivalent environments A. AVR setup. B. VVR setup. C. Metrically-equivalent and same-sensory tracks, illustrated in simplified diagrams. The three cues (1 to 3 in AVR1 and VVR1) and four cues (1 to 4 in AVR2 and VVR2) are labeled according to their identities, i.e., cue 1 in AVR1 and AVR2 was the same auditory cue at 15 kHz, and cue 1 in VVR1 and VVR2 was the visual cue with the same shape and pattern. Yellow indicates a hidden reward zone. D. The time the mice spent stopping in the four tracks in C. Gray indicates cue areas and yellow represents reward zone. E. Success rates in receiving reward. F. Number of active cells between pairs of metrically-equivalent tracks. p values: AVR1 versus VVR1: 0.0041; AVR2 versus VVR2: 0.0108. G. Percentage of common cells between track pairs. Magenta: AVRs (1: AVR1; 2: AVR2). Green: VVRs (1: VVR1; 2: VVR2). p values are shown in Table S1. H. Schematic of NN distance analysis. This example assumes that cells in groups A and B are anatomically separated, and A contained less cells than B. B was first randomly subsampled to be sample-matched with A. The distances between individual cells in A with their nearest neighbors (NN) in subsampled B (AB distances) were then calculated. The distances of cells in subsampled B to A cells (BA distances) were also calculated. The distances between groups A and B were the mean of AB and BA distances and were compared with shuffled distances, which were calculated for randomly permuted cells among A and subsampled B. If cells in A and B were anatomically separated, the distances between A and B should be longer than shuffled distances. I. An example FOV with active cells in AVR1 and VVR1, including unique cells in AVR1 (magenta) or VVR1 (green), and common cells of the two tracks (white). J. Cumulative distributions of distances between AVR1unique cells and VVR1 unique cells and shuffled distances. K-L. Similar to I-J but for active cells in AVR2 and VVR2. *p ≤ 0.05, **p ≤ 0.01, ***p ≤0.001, n.s. p > 0.05. Error bars represent mean ± SEM. Data was collected from 6 mice.

The mice were trained to navigate in all four tracks. In each pair of metrically-equivalent tracks, they specifically stopped at reward zones and received rewards at comparably high success rates (Figures 3D and 3E), indicating that both auditory and visual cues provided effective spatial information for navigation.

By tracking active MEC cells in the same set of FOVs in all four tracks, we found that VVRs recruited more cells than AVRs (Figure 3F). While the percentages of common cells in all track combinations were above random levels (Figure S2E), the percentages across metrically-equivalent tracks were lower than those across same-sensory tracks (within the same track and between different same-sensory tracks, Figure 3G and Table S1). This result indicates that cells recruited by metrically-equivalent tracks in different sensory modalities were more exclusive than those recruited by same-sensory tracks, supporting the existence of specific cell populations with different modality preferences.

We further investigated whether the cells uniquely active in metrically-equivalent tracks (e.g., AVR1 unique cells and VVR1 unique cells) were anatomically separated by comparing their anatomical distances with those in random cases^55^. We first identified the unique population with fewer cells (A) and calculated their distances with their nearest neighbors (NN) in the other unique population (B), which were randomly subsampled to be size-matched with A. The distances were then compared with shuffled distances, which were calculated by randomizing cells among A and subsampled B. If cells in A and B were anatomically separated, their distances would be longer than shuffled distances (Figure 3H). However, the distances between AVR1 unique cells and VVR1 unique cells were comparable with shuffled distances, indicating that they were not anatomically separated (Figures 3I and 3J). A similar conclusion was true for AVR2 unique cells and VVR2 unique cells (Figures 3K and 3L). These results indicate that unique cells in metrically-equivalent tracks were anatomically intermingled.

Thus, while auditory and visual cues provided metrically-equivalent spatial information to guide comparable navigation behaviors in unisensory tracks, they tended to recruit different cell populations that were anatomically intermixed in the MEC microcircuit.

### Features of unique and common cells in metrically-equivalent unisensory tracks support differential encoding of visual and auditory spatial information

We subsequently characterized the MEC map in metrically-equivalent tracks by examining calcium responses of MEC cells in each track. Since the MEC exhibited different responses in the correct and error runs in AVR1 (Figures 2O-Q and S1F-H), we first asked whether such a difference also existed in the other tracks. Indeed, the responses in AVR2, VVR1, and VVR2 all showed higher RBR consistency in the correct than in the error runs (Figures S2F-J). The consistency differences in AVRs were larger than those in VVRs (Figure S2F), suggesting that neural response in AVRs was more cognition-driven than in VVRs. Therefore, all following analyses only used the activity in the correct runs in all tracks.

Next, we compared the activity of unique cells in metrically-equivalent tracks. They contained “unimodality cells” that were only active in tracks of a specific sensory modality. Overall, unique cells in VVRs showed more robust responses than those in AVRs, as reflected by their higher calcium activity (Figure 4A), more spatial fields (Figures 4B-D), and higher RBR activity consistency (Figure 4E) compared to AVR unique cells. VVR unique cells also exhibited a higher ratio of field abundances in the areas within (in-cue) and outside of cues (out-cue) (Figure 4F), reflecting their higher activity specificity to cues compared to AVR unique cells. On the other hand, AVR unique cells preferentially represented the cues near reward, as reflected by the high ratio (>1) between the percentages of the cells with fields near and away from the reward (Figures 4G) and the higher activity consistency of the cells at cues near the reward than at those away from the reward (Figures 4H and 4I). However, such features were not observed in VVR unique cells.

**Figure 4:**
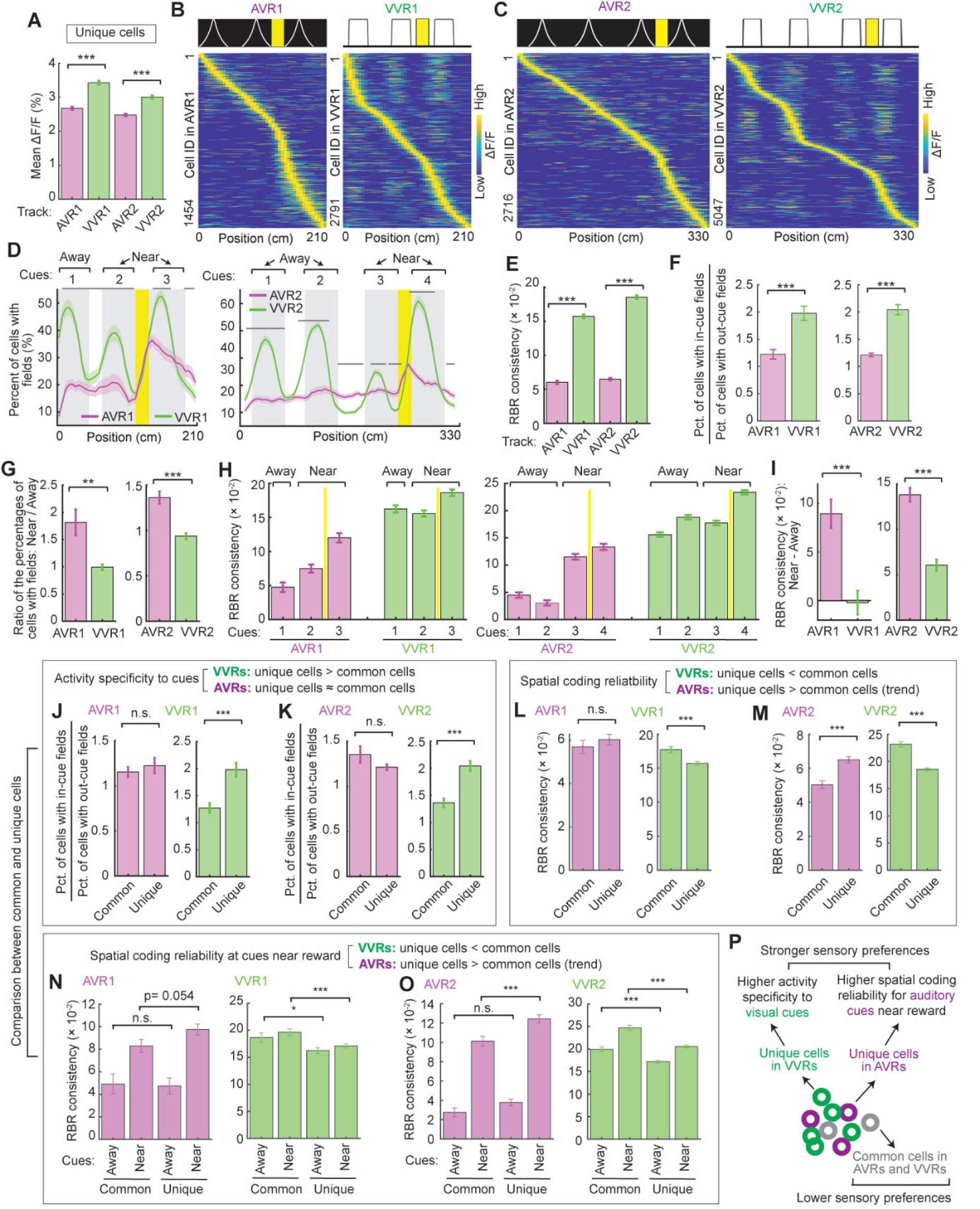
Unique and common cells in metrically-equivalent AVRs or VVRs A. Average calcium response (mean ΔF/F) of unique cells in metrically-equivalent AVRs or VVRs. p values: AVR1 versus VVR1: 1.6×10^−23^; AVR2 versus VVR2: 1.1×10^−19^. B and C. Activity matrices of unique cells in AVR1 or VVR1 (B), and in AVR2 or VVR (C). D. Percentage of unique cells with spatial fields in AVR1 versus VVR1 tracks (left) and in AVR2 versus VVR2 tracks (right). “Away” and “Near” indicate cues away from and near the reward. Same below. E. RBR activity consistency of unique cells. p values for AVR1 versus VVR1: 9.0×10^−113^; for AVR2 versus VVR2: 1.4×10^−233^. F. Ratio of the percentages of unique cells with in-cue and out-cue fields. p values for AVR1 versus VVR1: 5.5×10^−7^; AVR2 versus VVR2: 1.5×10^−11^. G. Ratio of the percentages of unique cells with fields at cues near and away from reward. p values for AVR1 versus VVR1: 0.0013; AVR2 versus VVR2: p = 1.1×10^−7^. H. RBR activity consistency in cue areas for unique cells in AVR1 or VVR1 (left) and in AVR2 or VVR2 (right). I. Normalized difference in RBR activity consistency in cue areas near and away from reward for unique cells in AVR1 or VVR1 (p = 2.3×10^−6^) and in AVR2 or VVR2 (p = 4.9×10^−15^). J - O. Comparison between common and unique cells. J. The comparison of the ratio of the percentages of cells with in-cue and out-cue fields for common and unique cells in AVR1 and VVR1(p = 6.4×10^−6^). K. Similar to J but for AVR2 and VVR2 (p = 2.8×10^−12^). L. The comparison of RBR activity consistency for common and unique cells in AVR1 and VVR1 (p = 3.3×10^−4^). M. Similar to L but for AVR2 (p = 1.8×10^−6^) and VVR2 (p = 1.5×10^−18^). N. The comparison of RBR activity consistency at cues near and away from reward for common and unique cells in AVR1 (p value for cues near reward is 0.054) and VVR1 (p values for cues near and away from reward are 2.9×10^−4^ and 0.0150, respectively). O. Similar to N but for AVR2 (p = 5.7×10^−4^ for cues near reward) and VVR2 (p values for cues near and away from reward are 4.5×10^−11^ and 1.3×10^−5^, respectively). P. Summary of activity features of unique and common cells in metrically-equivalent AVRs and VVRs. Unique AVR cells (magenta) and unique VVR cells (green) showed higher activity specificity to visual cues and higher spatial coding reliability for auditory cues near reward, respectively, indicating their stronger sensory preferences. Common cells showed weaker sensory preferences. *p ≤ 0.05, **p ≤ 0.01, ***p ≤0.001, n.s. p > 0.05. Error bars represent mean ± SEM. Data was collected from 6 mice.

We further explored the activity of common cells between metrically-equivalent tracks. They are “multimodality” cells because they were active in tracks in different sensory modalities. The above activity differences between unique cells were also observed in common cells (Figures S3A-I), except that common cells showed comparable activity specificities to the cues in AVRs and VVRs (Figures S3F). Thus, in both unique and common cells, VVRs were dominantly encoded over AVRs, whereas the cues near reward were more preferentially represented in AVRs than in VVRs, supporting different neural maps for the tracks in different sensory modalities. The dominant VVR encoding was not a consequence of faster running speed^56^, because the speeds in VVRs were similar or even slower than those in AVRs (Figure S3J).

Moreover, special features of unique cells distinguished them from common cells. In VVRs, the ratio of in-cue and out-cue field abundance was higher in unique cells than in common cells, indicating that VVR unique cells had higher activity specificity to visual cues (Figures 4J and 4K). This difference was not observed between unique and common cells in AVRs. In contrast, in AVRs, unique cells exhibited a trend of higher RBR activity consistency than common cells (Figures 4L and 4M), specifically at the cues near the reward (Figures 4N and 4O), indicating that AVR unique cells had higher spatial activity reliability to encode cues near reward. This feature was not observed in VVR unique cells, which showed lower activity consistency than common cells (Figures 4L-4O). These striking features of unique cells compared to common cells suggest that unique cells in VVRs and AVRs were specialized cell populations with stronger activity preferences to visual and auditory cues, respectively (Figures 4P).

In summary, metrically-equivalent tracks were differentially encoded in the MEC by both unique and common cell populations. VVRs were more robustly represented than AVRs, whereas AVR encoding was more biased toward cues around reward. Meanwhile, neural responses in same-sensory tracks shared more similarities. Moreover, unique cells in AVRs and VVRs showed stronger activity preferences to cues compared to common cells, supporting their “unimodality” feature for auditory and visual cues, respectively.

### Individual MEC cells have different and similar responses in metrically-equivalent and same-sensory tracks, respectively

While the above results at the population level indicated different responses of common cells in metrically-equivalent AVRs and VVRs (Figure S3A-I), we further investigated whether, at the individual cell level, the different responses were still observed, and whether common cells in same-sensory tracks exhibited greater similarities.

We first explored whether response patterns of the same cell remapped across metrically-equivalent tracks by calculating spatial activity correlations of individual common cells across the paired AVR and VVR tracks (Figure 5A). Activity correlations of the same cells in repeating sessions of AVR1 or VVR1 were higher than random correlations calculated by permuting activity pairs of individual cells in the repeating sessions, indicating that the same cell largely maintained similar activity patterns in the same track (Figures 5B and 5C, left two columns). In contrast, activity correlations of the same cells in AVR1 and VVR1 were below random (Figures 5B and 5C, right column), indicating activity remapping in the two tracks. Similar observations were made for AVR2 and VVR2 (Figures 5D and 5E). Therefore, the activity of the same cell remapped across metrically-equivalent tracks.

**Figure 5:**
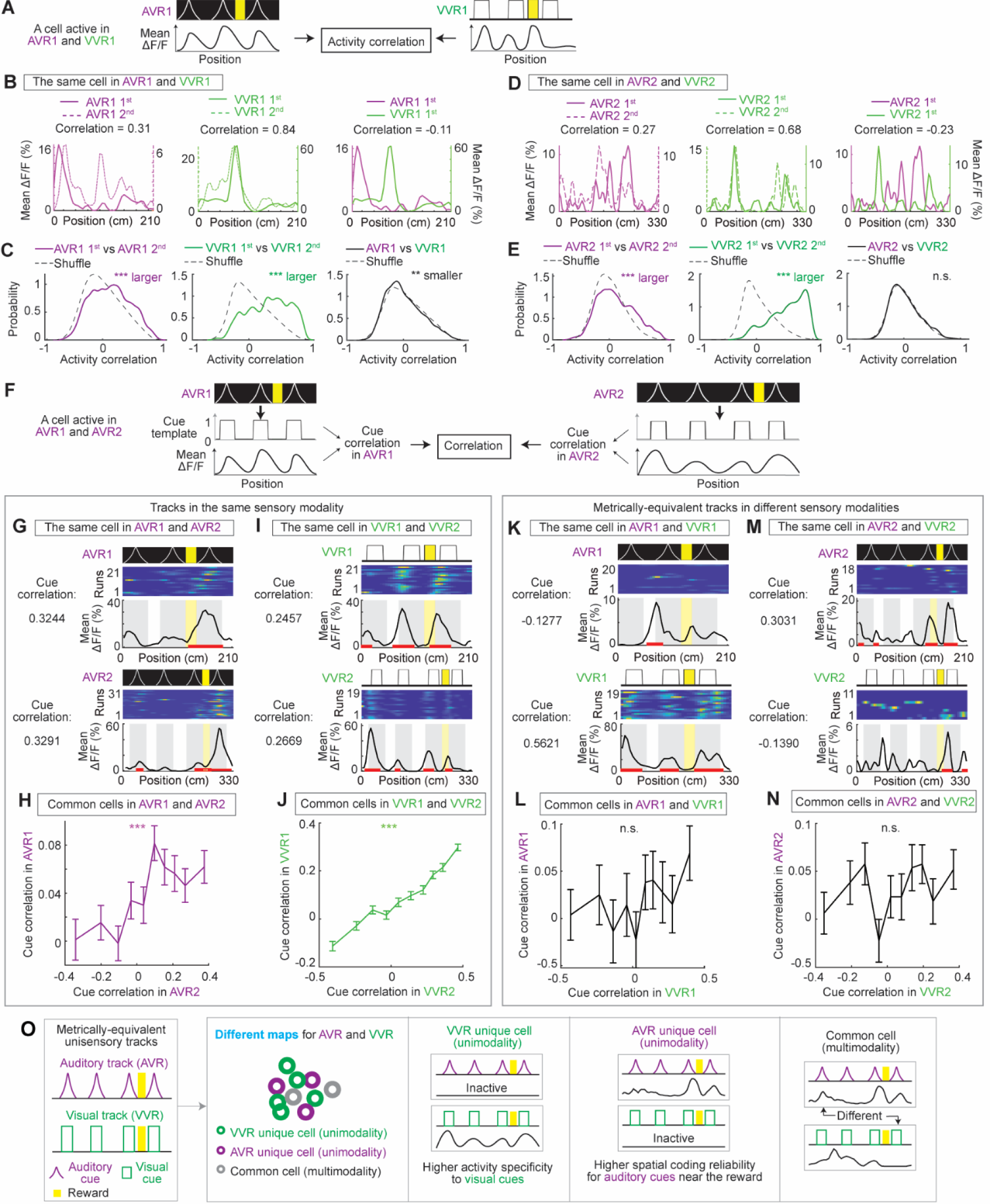
The activity of individual MEC cells in metrically-equivalent and same-sensory tracks. A. Diagram illustrating the calculation of activity correlation of a cell active in AVR1 and VVR1. B. Activity correlations of a cell in two sessions in AVR1 (left), in two sessions in VVR1 (middle), and in AVR1 and VVR1 (right). C. Distribution of activity correlations of common cells in two sessions of AVR1 (left), in two sessions of VVR1 (middle), and in AVR1 and VVR1 (right), and. All correlations are compared with shuffled distribution created by permuting activity pairs of individual cells in the two sessions. The texts next to asteroids indicate that real data were larger, smaller, or comparable (n.s.) to shuffles based on two-tailed Kolmogorov-Smirnov test. p values from left to right: 1.9×10^−51^ (real data are larger), 4.5×10^−89^ (real data are larger), and 0.0089 (real data are smaller). D and E. Similar to B and C, respectively, but for AVR2 and VVR2. For E: p values from left to right: 3.6×10^−13^, 9.4×10^−270^, and 0.0970. F. Diagram illustrating the calculation of cue correlations of a cell active in AVR1 and AVR2. G-N. Comparison of cue correlations of common cells in same-sensory tracks (G-J) and in metrically-equivalent tracks (K-N). G. A cell active in both AVR1 and AVR2 with its cue correlations in the two tracks (top). H. Significantly positive correlation of cue correlations of common cells in AVR1 and AVR2. P = 7.5×10^−5^. I-N. Similar to G and H but for cells active in VVR1 versus VVR2 (I and J, p = 1.1×10^−87^), AVR1 versus VVR1 (K and L, p = 0.1010), and AVR2 versus VVR2 (M and N, p = 0.2349). For H, J, L and N: Data was binned in every 10^th^ percentile of values on x-axis (e.g., cue correlation in AVR2 in H). O. Summary of encoding of metrically-equivalent AVRs and VVRs. Three types of cells were identified: unique cells in AVRs and unique cells in VVRs are unimodality cells with higher specificity to encode visual and auditory cues, respectively. Common cells had multimodality responses that differentially encoded AVRs and VVRs. Additionally, there were more unique cells in VVRs than in AVRs and more robust encoding of VVRs than AVRs. In contrast, the AVR encoding was more biased towards the most task-relevant cues near the reward. *p ≤ 0.05, **p ≤ 0.01, ***p ≤0.001, n.s. p > 0.05. Error bars represent mean ± SEM. Data was collected from 6 mice.

Although the remapping also occurs across different visual environments^57^, VVR tracks share a cue cell population that consistently encodes individual cues across different tracks^10^. While a significant population of shared cue cells were also identified between VVR1 and VVR2 (Figures S4A-C), no cue cells existed in AVRs (Figure S4D), likely because the MEC preferentially encoded auditory cues near reward, rather than all cues (Figure 2). Therefore, the MEC contains a dedicated visual, but not auditory, cue cell population.

Given the difference in cue cells in VVRs and AVRs, we assess the relationship between neural activity and cue arrangement based on “cue correlation”, which was the correlation between a cell activity and a “cue template” representing in-cue and out-cue areas of a track by ones and zeros, respectively (Figure 5F). We found that cue correlations were positively correlated for common cells in same-sensory tracks (Figures 5G and 5H for AVR1 and AVR2; Figures 5I and 5J for VVR1 and VVR2) but were uncorrelated in metrically-equivalent AVR and VVR tracks (Figures 5K and 5L for AVR1 and VVR1; Figures 5M and 5N for AVR2 and VVR2). These results imply that the cues in the same sensory modality shaped neural activity similarly even though they provided different metric information, whereas those in different sensory modalities shaped neural activity differently despite their equivalent metric information.

Lastly, we asked whether common cells in metrically-equivalent tracks exhibited general responses to reward locations independent of sensory modality, since a previous study identified modality-invariant hippocampal response to rewards^58^. We isolated “reward cells” as those with high “reward scores”, calculated as the normalized difference in calcium response within and outside of the reward zone (Figures S5A and S5B). We further identified “common reward cells” shared between metrically-equivalent tracks and compared their percentages on an individual-mouse basis with those at random levels (Figures S5C and S5D). Overall, no significant common reward cell populations were identified across metrically-equivalent tracks (Figures S5E-H), indicating that MEC reward response was also different in these tracks.

Overall, the responses of common cells (multimodality) in metrically-equivalent tracks completely remapped and showed different relationships with cue arrangements. In contrast, the relationship was similar for common cells in same-sensory tracks. These results indicate that cells differentially and similarly responded in environments in different and same sensory modalities, respectively. This conclusion and the above characterizations of unique cells (unimodality) in AVRs and VVRs strongly indicate that the MEC use different maps to represents spatial information in different sensory modalities (Figure 5O).

### Separate cells encoded visual and auditory cues in a multisensory track

Next, we explored the encoding of auditory and visual spatial information in a multisensory environment with both sensory cues (Figure 1B). The cues in AVR2 and VVR2 were arranged at nonoverlapping locations in a visual-auditory VR (VAVR) track (Figures 6A and S6A). To direct the mouse’s attention to both sensory cues, two reward zones were placed: one after an auditory cue and the other after a visual cue (auditory reward and visual reward, respectively), identical to those in AVR2 and VVR2, respectively (Figure S6A). Three of the above mice navigated in VAVR track. They specifically stopped in both reward zones (Figure 6B) and received both rewards at comparable success rates. In ∼ 40% of the runs, mice correctly identified both rewards within the same run (Figure 6C). Hence, the mice could use both auditory and visual cues to locate rewards without sensory bias.

**Figure 6:**
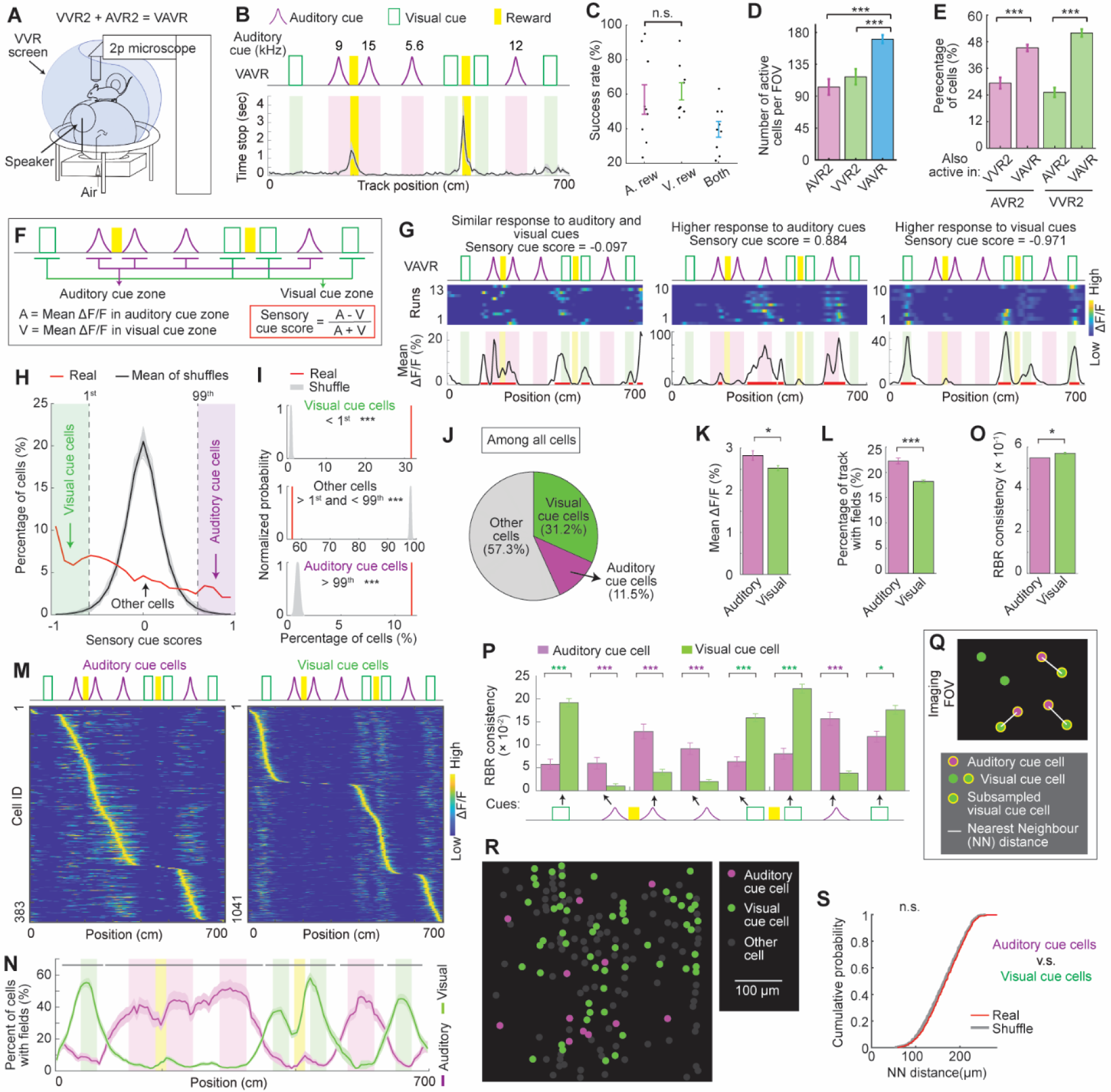
Separate encoding of auditory and visual cues in VAVR A. VAVR and imaging setup. The use of two, instead of eight, speakers for auditory cues was due to space limitation within the screen. B. The time the mice spent stopping along VAVR track. C. Success rates in receiving reward following an auditory cue (A. rew), a visual cue (V. rew), and receiving both rewards in the same run (Both). D. Numbers of active cells per FOV in AVR2, VVR2, and VAVR. p values for AVR2 versus VAVR: 1.0×10^−6^; VVR2 versus VAVR: 4.1×10^−5^. E. The percentage of AVR2 cells that were active in VVR2 (bar 1 from the left) and VAVR (bar 2), and the percentage of VVR2 cells that were active in AVR2 (bar 3) and VAVR (bar 4). p values for VVR2 versus VAVR: 2.4×10^−7^; AVR2 versus VAVR: 4.3×10^−15^. F. Calculation of sensory cue score. G. Examples of cells responding similarly to both auditory and visual cues (left), stronger to auditory cues (middle), and stronger to visual cues (right), and their sensory cue scores. H. Distributions of real and shuffled sensory cue scores (gray: 100 shuffled datasets; black: mean). Vertical dashed lines indicate the 1^st^ and 99^th^ percentile of shuffle used as thresholds for identifying visual (in green zone) and auditory cue cells (in magenta zone), respectively. I. Comparisons between real (red) and shuffled cases (gray) in the percentages of cells with sensory cue scores below 1^st^ threshold (p_above = 0), between 1^st^ and 99^th^ thresholds (p_below = 0), and above 99^th^ threshold (p_above = 0). Similar to Figure S5, p_above and p_below indicate the percentage of shuffled datasets, in which the percentage of a certain cell type (e.g., those above 1^st^ threshold) was above or below the percentage of the real dataset. J. Percentages of visual, auditory cue cells, and other cells among all active cells in VAVR. K. Average calcium response (mean ΔF/F) of visual and auditory cue cells. p = 0.0143. L. Percentage of track with spatial fields of auditory and visual cue cells. p = 3.3×10^−12^. M. Activity matrices of auditory and visual cue cells. N. Percentage of auditory and visual cue cells with spatial fields along VAVR track. O. RBR activity consistency of auditory and visual cue cells. p = 0.0301. P. RBR activity consistency of auditory and visual cue cells at individual cues. Significant p values (from left to right): 1.0×10^−4^; 1.8×10^−5^; 2.2×10^−7^; 1.9×10^−11^; 1.3×10^−8^; 6.3×10^−6^; 1.7×10^−8^; 0.0217. Q. Schematic of NN distance analysis for auditory and visual cue cells, similar to Figure 3H. R. An example FOV with auditory cue cells, visual cue cells, and other cells. S. Cumulative distributions of NN distances between auditory and visual cue cells and shuffle distances. *p ≤ 0.05, **p ≤ 0.01, ***p ≤0.001, n.s. p > 0.05. Error bars represent mean ± SEM. Data was collected from 3 mice.

MEC dynamics of the same set of FOVs in AVR2 and VVR2 were imaged during VAVR navigation. Interestingly, VAVR activated more cells than AVR2 and VVR2 (Figure 6D). A higher percentage of AVR2 cells was active in VAVR than in VVR2 (Figure 6E, the left two columns), indicating that VAVR activated AVR2 unique cells relative to VVR2. Similarly, more VVR2 cells were active in VAVR than in AVR2 (Figure 6E, the right two columns). Thus, combining auditory and visual cues in VAVR recruited unique cells in AVR2 or VVR2, consistent with the above observation that these unique cells had strong sensory preferences (Figure 5O). They were therefore recruited when cue in both sensory modalities were combined in VAVR.

To further explore the possibility that visual and auditory spatial information in VAVR is encoded by separate cells (Figure 1B, b), we investigated cell populations specific for auditory and visual cues in VAVR based on “sensory cue scores”, which represented normalized difference in individual cell responses in auditory and visual cue zones (Figure 6F). The cells with sensory cue scores around zero had comparable responses to both types of cues, whereas those with scores close to 1 and −1 had higher responses to auditory and visual cues, respectively (Figure 6G). If visual and auditory cues were unbiasedly represented by the cell population, sensory cue scores should exhibit a Gaussian distribution centered around zero, which was true for shuffled cue scores calculated from randomly permuted calcium responses of individual cells (Figure 6H, black curve). However, sensory cue score distribution of real cells was highly skewed to lower values, indicating an overall stronger response to visual cues at the individual cell level (Figure 6H, red curve).

Using the 1^st^ and 99^th^ percentiles of the shuffled scores as thresholds, we identified “visual cue cells” and “auditory cue cells” with relatively higher responses to visual and auditory cues, respectively (Figure 6H). The percentages of these cells were higher than those expected by chance, and the percentage of other cells between the thresholds was lower (Figure 6I), reflecting robust modality-specific encoding by these cells. To confirm that the encoding was due to the different sensory modalities of the cues, rather than their different track locations, we compared the distribution of real sensory scores with a set of “fake cue scores”, which were calculated by randomly assigning the eight cues into four fake auditory cues and four fake visual cues (Figure S6B). In comparison to “fake” cases, the percentages of real auditory and visual cue cells were higher, and the percentage of real other cells was lower (Figure S6C), indicating that the different sensory modalities of the cues indeed led to their specific encoding.

Calcium responses of visual and auditory cue cells further supported their sensory specificity. More visual cue cells were identified than auditory cue cells (Figure 6J), whereas auditory cue cells had higher calcium responses (Figure 6K) and higher spatial field prevalence on the track than visual cue cells (Figure 6L). The fields of auditory and visual cue cells specifically distributed around their preferred cues (Figures 6M and 6N). Visual cue cells had higher RBR activity consistency than auditory cue cells (Figure 6O), but on an individual-cue basis, the activity of auditory and visual cue cells were more spatially consistent at their preferred cues (Figure 6P). Anatomically, auditory and visual cue cells were intermingled, as indicated by their anatomical distances that were comparable to shuffle distances (Figure 6Q-S).

Moreover, we asked whether auditory and visual cue cell ensembles varied in different behavioral conditions, in which the mice received both rewards (Both, used in above analyses), only auditory rewards (Auditory), only visual rewards (Visual), and no reward (None) (Figure S7A). We found that the fractions of auditory cue cells, visual cue cells, and other cells were largely similar across the four behavioral conditions, except for a slightly higher fraction of auditory cue cells in “Auditory” runs (∼5% increase) (Figures S7B-E), consistent with the stronger cognitive modulation of auditory response (Figure S2F). For the three cell types, significant overlaps were observed between the cells in “Both” and in the other behavioral conditions (Figure S7F), further indicating the largely stable cell compositions. Interestingly, the cells showed differences in their RBR activity consistencies along the track according to behaviors. In comparison to the “Both” condition, auditory cue cells exhibited reduced consistency when auditory reward was not received (in “Visual” and “None” conditions) (Figure S7G), whereas visual cue cells had lower consistency when none of the rewards was received (in the “None” condition) (Figure S7H). These results indicate that spatial activity consistency, rather than sensory preference of the cells, varied with behaviors.

In summary, the existence of separate cell populations responding to auditory and visual cues indicates differential encoding of these two types of sensory cues.

### Separate MEC cells responded to either visual or auditory rewards in a multisensory environment

Next, we investigated neural responses within visual and auditory reward zones in VAVR. While modality-invariant reward responses were absent in unisensory metrically-equivalent tracks (Figure S5), we asked whether such responses existed in VAVR when mice successfully localized both auditory and visual rewards in the same run. Similar to the approach used in unisensory tracks (Figure S5A), reward cells in VAVR were classified as those with high “reward scores” (Figures 7A and 7B). The reward zone combined both auditory and visual reward zones to unbiasedly identify all cells responding to either or both zones. However, strikingly, classified reward cells exhibited robust spatial fields in either zone, implying their strongly biased encoding of reward associated with cues in a specific sensory modality (Figure 7C). To reveal the difference in reward cell responses in auditory and visual reward zones, we calculated “sensory reward scores” (Figure 7D). Indeed, sensory reward scores exhibited a strong bimodal distribution (Figure 7E), indicating specific encoding of either auditory or visual reward zones.

**Figure 7:**
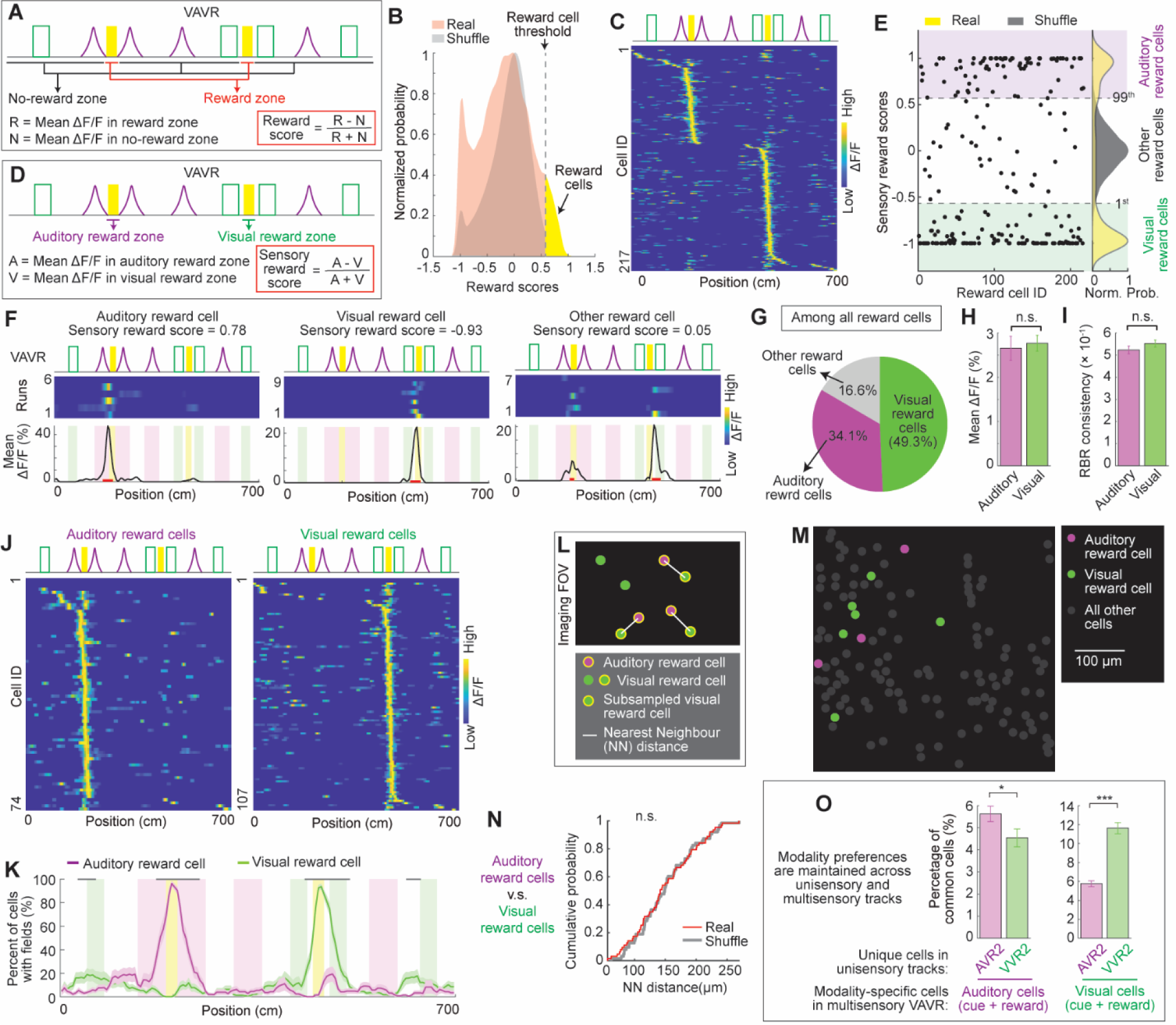
Separate encoding of visual and auditory rewards in VAVR. A. Calculation of reward score. B. Distribution of real reward scores in real cells and shuffles. The vertical dash line indicates the reward cell threshold, which is the 99^th^ percentile of shuffles. The yellow area is classified reward cells above the threshold. C. Activity matrix of reward cells. D. Calculation of sensory reward scores. E. Distribution of sensory reward scores of classified reward cells in B. Left: each dot indicates the sensory reward score of one reward cell. Right: the distributions of sensory reward scores (yellow) versus shuffled scores (gray). Dashed gray lines are the thresholds at 1^st^ and 99^th^ percentiles of shuffles to classify visual and auditory reward cells, respectively. The middle cells are other reward cells. F. Examples of auditory, visual, and other reward cells. G. Fractions of auditory, visual, and other reward cells among all reward cells. H. Average calcium response (mean ΔF/F) of visual and auditory reward cells. I. RBR activity consistency of auditory and visual reward cells. J. Activity matrices of auditory and visual reward cells. K. Percentage of auditory and visual reward cells with spatial fields along VAVR track. L. Schematic of NN distance analysis for auditory and visual reward cells, similar to Figure 3H. M. An example FOV with auditory, visual reward cells, and all other cells. N. Cumulative distributions of NN distances between auditory and visual reward cells and NN distances in shuffles. O. The percentage of shared cells between unique cells in unisensory AVR2 or VVR2 and modality-specific cells in VAVR. Visual and auditory cells in VAVR included the corresponding cue and reward cells. p values are 0.0486 (left) and 8.6×10^−13^ (right). * p/p_above ≤ 0.05, ** p/p_above ≤ 0.01, *** p/p_above ≤0.001, n.s. p/p_above > 0.05. Error bars represent mean ± SEM. Data was collected from 3 mice.

We further classified visual and auditory reward cells as those with sensory reward scores below and above the 1^st^ and 99^th^ percentiles of shuffled scores (calculated after randomizing reward cell responses in auditory and visual reward zones), respectively (Figure 7E, right). Most reward cells fell into these two categories. Other reward cells with sensory reward scores between the thresholds constituted a smaller fraction (Figures 7F and 7G). Auditory and visual reward cells showed similar calcium responses (Figure 7H) and RBR activity consistencies (Figure 7I). They largely exhibited spatial fields at their preferred reward zones (Figures 7J and 7K). They were anatomically intermingled according to their comparable distances with shuffles (Figures 7L-N).

These results further demonstrated that reward response in the MEC is modality-specific. Although the existence of the reward is a type of cognitive information, when rewards were associated with cues in different sensory modalities, they were represented by separate cells, highlighting sensory information as an important component of the MEC map.

Given the identification of unimodality cells in several environments, including unique cells in AVR2s or VVR2, and visual and auditory cells (cue cells and reward cells) in VAVR, we further asked whether these cells maintained their sensory preferences across unisensory and multisensory environments. Interestingly, auditory cells in VAVR shared a higher percentage of common cells with AVR2 unique cells than with VVR2 unique cells. Similarly, visual cells in VAVR shared a higher percentage of common cells with VVR2 unique cells than with AVR2 unique cells (Figure 7O). These results imply that modality-specific cells were largely consistent across environments, rather than being randomly assigned in each environment.

### Multimodality MEC cells differentially responded to visual and auditory information in the multisensory environment

Lastly, we examined the activity of cells that did not belong to the above auditory and visual cells (cue and reward cells). These cells had relatively lower sensory bias, and therefore, were considered “multimodality” cells for both visual and auditory spatial information. Multimodality cells constituted 54.6% of the total cells (Figure 8A) and exhibited spatial fields spanning the entire VAVR track (Figure 8B). At the population level, they showed higher calcium responses (Figure 8C) and more spatial fields (Figure 8D) around visual cues than around auditory cues. Their activity was also more spatially consistent near visual cues and visual reward (Figure 8E). We further compared individual cell responses within auditory and visual information zones, which included cues and reward (Figure 8F). As expected, individual cells showed higher calcium response (Figure 8G), wider coverage of spatial fields (Figure 8H), and higher RBR activity consistency (Figure 8I) within visual information zone than within auditory information zone. These results revealed preferential encoding of visual spatial information by multimodality cells.

**Figure 8.**
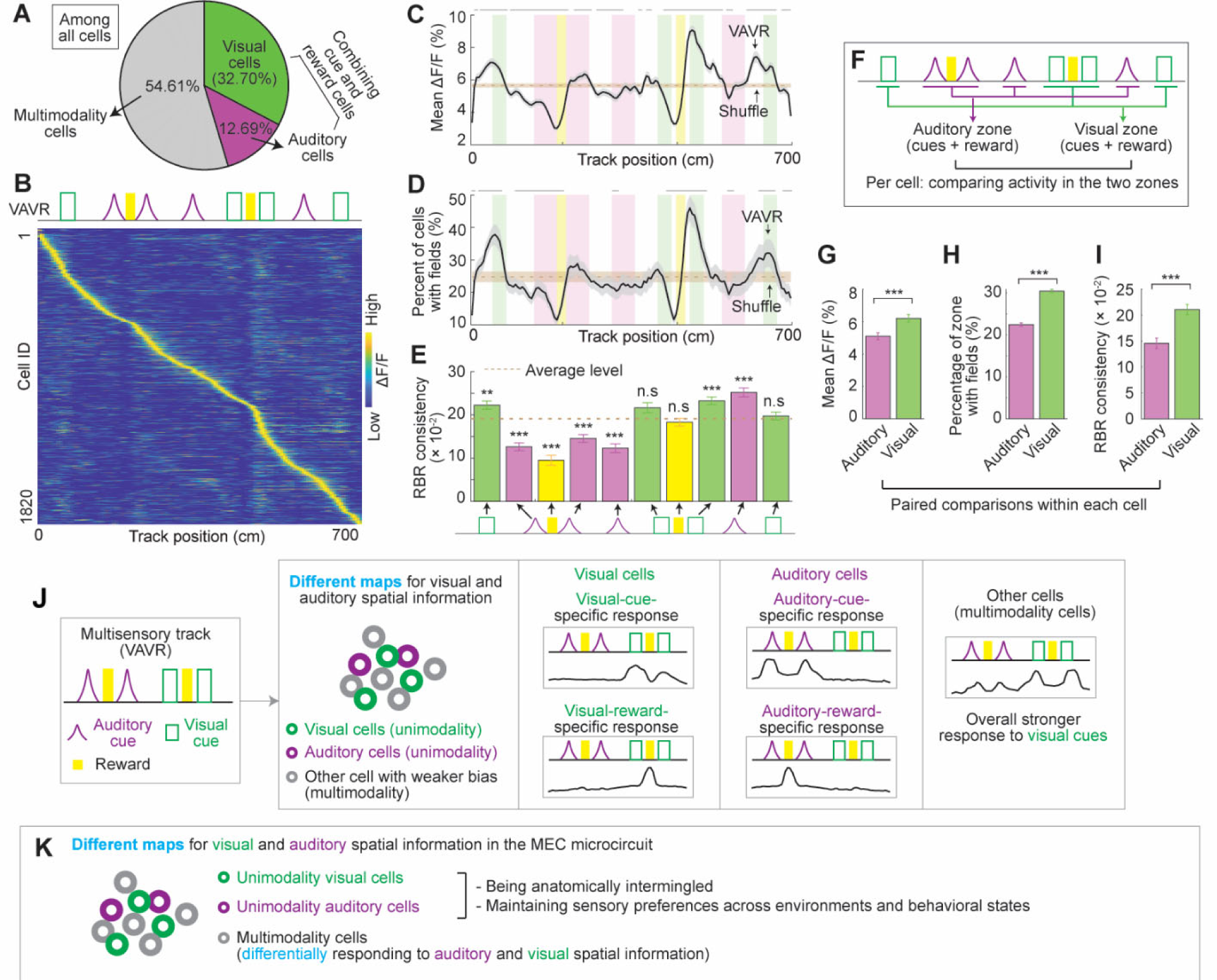
Calcium response of multisensory cells in VAVR and summary diagrams A. The percentage of visual, auditory cells (including cue cells and reward cells), and multimodality cells. B. Activity matrix of multisensory cells. C. Calcium response amplitude (mean ΔF/F) of multimodality cells across the track. D. Percentage of multimodality cells with spatial fields along VAVR track. E. RBR activity consistency of multimodality cells at individual cues and rewards. Significant p values from left to right: 0.0038, 1.0×10^−9^, 3.1×10^−11^, 2.5×10^−5^, 1.7×10^−6^, 2.7×10^−5^, 1.5×10^−5^. F. A diagram showing the comparison of activity features of individual multimodality cells in auditory and visual zones, which include cue and reward areas. G. Comparison of averaged calcium response (mean ΔF/F) in auditory and visual zones on individual-cell basis. p value for paired t test: 9.6×10^−146^. H. Comparison of field coverage in auditory and visual zones (percentage of zones with fields) on individual-cell basis. p value for paired t test: 2.5×10^−50^. I. Comparison of RBR activity consistency in auditory and visual zones on individual-cell basis. p value for paired t test: 2.5×10^−8^. J. Summary of multisensory encoding in VAVR. Three types of cells were identified in the MEC: visual cells and auditory cells were unimodality cells specific for visual and auditory spatial information (cue and reward), respectively. Other cells had multimodality responses, which showed weaker sensory biases but still more strongly represented visual cues. Additionally, there were more visual cells. K. Overall summary based on the results in unisensory and multisensory tracks. The MEC uses different maps for auditory and visual spatial information. The microcircuit contained unimodality auditory cells and unimodality visual cells with high sensory preferences. There are also multimodality cells with lower sensory preferences, but they still exhibit different responses to auditory and visual information. Unimodality cells are anatomically intermingled and maintain their sensory preferences across multiple environments. *p ≤ 0.05, **p ≤ 0.01, ***p ≤0.001, n.s. p > 0.05. Error bars represent mean ± SEM. Data was collected from 3 mice.

Overall, the results in the multisensory environment led to similar conclusions as those in the unisensory environments that spatial information in visual and auditory modalities was specifically encoded by unimodality cells (visual and auditory cells) and differently encoded by multimodality cells, supporting different maps for spatial information in different sensory modalities (Figure 8J).

## Discussion

Here we uncovered the neural mechanism of multisensory encoding in the MEC. We found that while both visual and auditory spatial cues guided comparable reward-seeking behaviors in mice, the MEC circuit displayed different maps for spatial information in the two sensory modalities. The different maps were consistently revealed in two conditions: metrically-equivalent unisensory tracks (AVRs and VVRs) (Figure. 5O) and a multisensory track (VAVR) (Figure. 8J). In the unisensory condition, metrically-equivalent VVR and AVR tracks recruited unique cells with more specific encoding of sensory cues compared to common cells across the tracks. Common cells exhibited different population activity patterns in VVRs and AVRs, and at the individual cell level, showed complete activity remapping and distinct activity relationships with cue arrangement on the tracks. In the multisensory condition, visual and auditory spatial information, including cues and their associated rewards, was represented by separate cell ensembles. The remaining cells preferentially encoded visual spatial information.

Conversely, spatial information in the same sensory modality was encoded by the same map. In the unisensory condition, same-sensory tracks shared a higher percentage of common cells than those in different sensory modalities. MEC cells exhibited similar population responses in same-sensory tracks and common cells between the tracks showed consistent activity relationships with cue arrangement. In the multisensory condition, unique cells in AVR or VVR were recruited in VAVR, and auditory and visual cells in VAVR had higher overlaps with unique cells in AVR and VVR, respectively, indicating consistent modality preferences of the cells across unisensory and multisensory tracks.

Thus, the difference between visual and auditory encoding and the similarity between same-sensory encoding strongly indicate that the MEC forms specific and stable maps for auditory and visual spatial information (Figure. 8K). The maps were constructed by three cell populations: unimodality visual cells and unimodality auditory cells with higher sensory preferences to visual and auditory spatial information, respectively, and multimodality cells with lower preference but different responses to the two types of sensory information. Visual and auditory cells largely maintained their sensory preferences across tracks and were anatomically intermingled at the microcircuit level. Therefore, the MEC cognitive map not only encodes metric, but also sensory information of space. This multisensory capacity potentially allows the MEC to precisely encode multisensory information in nature.

### Modality-specific response in the MEC

In AVRs, the specific MEC response to the most task-relevant cues around reward during successful navigation indicates that the response was mainly cognition-driven, consistent with the anatomy that the MEC mostly receives indirect and processed auditory information^41–46,48^. In contrast, although MEC response in VVRs was also affected by cognitive states, the effect was weaker than that in AVRs (i.e., correct and error runs in Figure S2F, and different run conditions in Figure S7). In addition, VVRs recruit cue cells, which encode individual contralateral cues in multiple tracks, and therefore, are potentially driven by visual inputs^10^ (Figures S4A-C). These results suggests that MEC response to visual cues contains both cognitive and sensory components, consistent with the indirect and direct inputs from primary and secondary visual cortex to the MEC^42,45,49–52^. Therefore, we propose that different patterns of visual and auditory inputs to the MEC contribute to the differential encoding of visual and auditory spatial information. The more robust visual inputs could underlie the dominant encoding of visual over auditory cues in unisensory and multisensory environments (Figures 4, S3 and 8).

Notably, although the MEC differentially mapped visual and auditory cues, both types of cues guided comparably successful navigations in metrically-equivalent unisensory tracks (Figure 3E) and the multisensory track (Figure 6C), indicating that similar goal-directed navigation can be supported by very different cognitive maps. This observation is analogous to an earlier finding, in which Egyptian fruit bats navigated to the same goals in a room using either vision or echolocation, but their hippocampus exhibited completely different maps in the two sensory conditions^59^. Although the study hypothesized that the MEC could align metric information in different sensory maps into a modality-invariant cognitive map^59^, our observations of modality-specific encoding in the MEC suggest that this is unlikely the case.

As mentioned above, a modality-invariant response for reward was identified in the hippocampus when mice learned to associate reward with the first cue on the track regardless of visual or olfactory nature of the cues, suggesting that the cognitive map can represent abstract metric information when it must generalize the information across different sensory modalities^58^. However, such responses were not observed in the MEC, which did not show general reward responses across metrically-equivalent AVRs and VVRs (Figure. S5) and exhibited strongly biased activity to either auditory or visual reward zones in VAVR track (Figure 7). These results indicate that the MEC contains modality-variant reward response. Spatial cues in different sensory modalities likely provided different contexts around the reward, and therefore, led to differential encoding of reward zones, further emphasizing the importance of sensory information in shaping the MEC map. This observation is consistent with a recent study, in which the MEC primarily displays context-specific codes, rather than ongoing navigational goals per se^60^.

### Consistent MEC response to the same sensory modality

We found that unisensory tracks in the same sensory modality were similarly encoded by the MEC. AVRs consistently recruited smaller numbers of active cells than VVRs (Figure 3F). The tracks in the same sensory modality but with different metric information similarly shaped MEC population activity patterns (Figure 4 and S3) and shared higher percentage of common cells (Figure 3G), which showed consistent activity relationships with cue arrangements (Figures 5G-J). Moreover, in comparison to unisensory AVR2 and VVR2 tracks, multisensory VAVR activated the largest number of cells, including those uniquely active in AVR2 or VVR2, strongly implying the recruitment of unimodality cells when cues in both modalities were combined in the same environment (Figure 6E). Unique cells in a unisensory track were more likely to encode spatial information in the same sensory modality in VAVR (Figure 7O). In addition, the compositions of visual and auditory cue cells in VAVR were independent of behaviors (Figure S7). These results suggest that the MEC contains dedicated and separated cell populations for visual cues and auditory cues, and sensory preferences of the cells were largely stable across different environments and behavioral states. In contrast, sensory responses of hippocampal cells were more flexibly altered with contexts and behaviors^37^. Individual hippocampal cells could respond to the same odor^39^ or sound cue^38^ in one environment but not in another. In a multisensory track where the navigation goal was associated with either visual or olfactory cues, the total number of hippocampal neurons was preserved but the fractions of neurons with different modality preferences greatly altered according to the different goals^58^. Interestingly, it was recently reported that in the same task, the MEC predominantly encoded visual cues regardless of the different goals^61^, consistent with our observation that sensory preference of MEC cells is independent of behavior. Thus, it is likely that separate MEC cells receive different inputs that shape their stable sensory preferences, whereas hippocampal cells receives integrated sensory inputs that drive their flexible sensory preferences^33^.

However, if the MEC indeed allocates a dedicated cell population for each sensory modality, the network would eventually run out of cells for various types of information. Therefore, the maps for certain sensory modalities must show greater overlaps than those for visual and auditory modalities. It is possible that the information in several sensory modalities with converged inputs to the MEC share a more similar map, such as somatosensory and auditory modalities, both of which indirectly arrive at the MEC through the PHC/RSC pathway^42,62^. It is also possible that the robust map for visual modality is different from all other sensory modalities because visual cortices provide the only direct sensory inputs to the MEC based on the current knowledge^42,45,49–52^. Additional studies on the MEC encoding of other sensory modalities and the anatomy of other sensory projections to the MEC are needed to test these possibilities.

In summary, modality-specific responses in the MEC in our study strongly support the representation of both metric and sensory information in the cognitive map. This feature likely enables accurate spatial encoding in multisensory environments, facilitates adaptable navigation using alternative sensory modalities, and underlies impaired multisensory navigation during aging.

## Acknowledgements

We thank all colleagues in the Gu laboratory for supporting the work, the Section on Instrumentation at National Institute of Mental Health for helping with building the virtual reality setups, the Mouse Auditory Testing Core at NIDCD for help with testing mice hearing and advice on task design. We also thank Dr. Ling-Gang Wu and Dr. Lorna Role for constructive comments on the manuscript.

## Author contributions

Conceptualization: DN, YG

Methodology: DN, YG

Investigation: DN, YG, GW

Visualization: DN, YG

Funding acquisition: YG

Supervision: YG

Writing – original draft: DN, YG

Writing – review & editing: DN, YG, GW

### Data and materials availability

Data and custom MATLAB codes will be available upon reasonable request to the corresponding author.

## Supplementary Materials

Materials and Methods

Figures S1 to S8

Table S1

## Materials and Methods

### Animals

All animal procedures were performed in accordance with animal protocol 1524 approved by the Institutional Animal Care and Use Committee (IACUC) at NIH/NINDS. For two-photon imaging experiments, GP5.3 mice (C57BL/6J-Tg (Thy1-GCaMP6f) GP5.3Dkim/J, JAX stock #028280) were used ^53^. These included 3 males and 4 females ranging from 3-4 months old at the time when first imaging began. Mice were maintained on a reverse 12-hr on/off light schedule with all experiments being performed in the light off period.

### Microprism Construction

Microprism construction procedures were similar to those described previously^25,54^. A canula (MicroGroup, 304H11XX) was attached to a circular cover glass (3mm, Warner Instruments, 64-0720). A right angle microprism coated with aluminum on the hypotenuse (1.5mm, OptoSigma, RBP3−1.5-8-550), was then attached to the opposite side of the cover glass. All attachments were performed using UV-curing optical adhesive (ThorLabs, NOA81).

### Microprism Implantation Surgery

Microprism implantation procedures were similar to those described previously^25,54^. Mice were anesthetized using a tabletop laboratory animal anesthesia system (induction: 3% isoflurane, 1L/min oxygen, maintenance: 0.5%−1.5% isoflurane, 0.7L/min oxygen, VetEquip, 901806) and surgery was performed on a stereotaxic alignment system (Kopf Instruments, 1900). A homeothermic pad and monitoring system (Harvard Apparatus, 50-7220F) was used to maintain a body temperature of 37°C. After anesthesia induction, dexamethasone (2mg/kg, VetOne, 13985-533-03) and saline (500µL, 0.9% NaCl, McKesson, 0409-4888-50) were administered by intraperitoneal (IP) injection, and slow-release buprenorphine (1mg/kg, ZooPharm, Buprenorphine SR-LAB) was administered subcutaneously. Enroflox 100 (10mg/mL, VetOne, 13985-948−10) was used as an antimicrobial wash just after the skull was exposed and just prior to sealing the skull. All insertions were performed on the left hemisphere, aligning with previous observations of more favorable vasculature on the left side^25^. A 3mm craniotomy was performed centered at 3.4mm lateral to the midline and 0.75mm posterior to the center of the transverse sinus (approximately 5.4mm posterior to the bregma). A durotomy was then performed over the cerebellum. Mannitol (3g/kg, Millipore Sigma, 63559) was administered by IP prior to the durotomy. The microprism was inserted into the transverse sinus and sealed to the skull with Vetbond (3M, 1469SB). The head plate was then mounted on the skull opposite the craniotomy. Finally, the prism and head plate were adhered to the skull with Metabond (Parkell).

### Visual Virtual Reality Setup

For all visual-behavioral experiments, a customized virtual reality (VR) setup was used, which projects a one-dimensional (1D) virtual environment based on the running of a mouse, similar to that described previously^54^. Mice were head-fixed onto an air-supported polystyrene ball (8” diameter, Smoothfoam) using the mounted head plate. The ball rotated on an axle, allowing only forward and backward rotation. The virtual environment was projected onto a dome screen filling the visual field of mice (270° projection). An optical flow sensor (Paialu, paiModule_10210) with infrared LEDs (DigiKey, 365−1056-ND) was used to measure the rotation of the ball and thereby control the motion of the virtual environment. The optical flow sensor output to an Arduino board (Newark, A000062), which transduced the motion signal to the computer controlling the virtual reality. An approximately 4μl water reward was provided via a lick tube at fixed locations (1 or 2 reward locations) in a given environment using a solenoid. A lick sensor connected to both the lick tube and head plate holder was used to detect mouse licking. A mouse licking the lick tube created a closed circuit between the lick sensor, the lick tube, the mouse (from the tongue to the skull), the headplate (which directly contacts the skull), and the head plate holder. The solenoid and lick sensor were controlled using a Multifunction I/O DAQ (National Instruments, PCI-6229). The virtual environments were generated and projected using ViRMEn software^20^. Imaging and behavior data were synchronized by recording a voltage signal of behavioral parameters from the VR system using the DAQ. ViRMEn environments were updated at 60Hz. The DAQ input/output rate was 1kHz. The synchronization voltage signal was updated at 20kHz. Final behavioral outputs were matched to the imaging frame rate (30Hz, see two-photon Imaging) for synchronization.

Environments were projected through a wratten filter (Kodak, 53-700) to reduce contamination of the imaging path with projected light. Virtual environments were 1D linear tracks with patterned walls and patterned visual cues at fixed locations. At the end of the track, mice were immediately teleported to the start of the track. Imaging experiments used 210cm (VVR1) and 330cm (VVR2) tracks.

### Auditory Virtual Reality Setup

For auditory-behavioral experiments, the same fundamental setup for visual VR was used in auditory VR for basic functions including motion signal detection, water delivery, and lick detection with no visual projection of the VR. The delivery of auditory cues as a function of 1D virtual track locations was achieved by a customized auditory-virtual-reality (AVR), which played continuous virtual sounds around a head-fixed mice (Figure 2A) running in darkness. Sounds were generated using a system of 8 speakers (SB acoustic SB21RDC) placed in a circular position with mouse head being the center (Figure S1A). The sound sources generated pure tones of frequencies 9, 12 and 15 kHz (for 210 cm AVR1 track) and 5.6, 9, 12, 15 kHz (for 330 cm AVR2 and 700 cm VAVR tracks), with a peak intensity at 94 dB SPL (sound pressure level in decibels with respect to 20 μPa, measured using a calibrated microphone, Earthworks Audio M50, calibrated by manufacturer). Sounds reached the peak intensity when mice were at the center of the audio zones in the AVR, intensity decreased by 6 dB when the virtual distance between center and mice doubled, decreased to ∼65 dB which is roughly equal to the room’s background noise. Spatial orientation effect of a mouse running toward or away from an auditory cue was mimicked by sequentially delivering the cue from the front to the back speakers, as described previously^17^.

Sound intensity was controlled using a second MATLAB interface with PsychToolbox software (PsychPortAudio library) which received positional information input from the first MATLAB interface that controlled ViRMEn software. Speaker systems were controlled by the specialized sound card (Audioscience ASI5780) that allows control of multiple speakers at the same time independently.

### Combining Auditory and Visual Virtual Reality

For combined-auditory-visual-behavioral (VAVR) experiments, the same setup for visual VR was used with additional two speakers positioned to the left and right sides of mice’s head within the projection dome (Figure 6A). The scheme to control each modality is described in above sections. Since there was not enough space inside the dome for all 8 speakers in AVR, we sacrificed the ability to simulate sound orientation, and the only information mice received from auditory cues was the distance from the cue center, which was achieved by varying the intensity of individual auditory cues. The VAVR track was 700 cm long and used the same set of cues in AVR2 and VVR2.

### Behavior training and imaging timeline

#### Early training to navigate in VR

Mice were allowed to recover for 5 days post-surgery and were then placed under water restriction, receiving 1ml water per day. After approximately 3 days of water restriction, mice were trained daily, starting from visual VR. Mice were trained to run in a 10 m 1D track for 1-4 weeks until mice began running consistently (> 30 runs per 40-minute training session for multiple consecutive days). Mice then were switched to auditory VR for auditory-behavioral training.

#### Behavioral training in AVR1

Mice started from a 210 cm AVR track (AVR1) and in passive task, during which mice only needed to run to a certain location in AVR to retrieve water. After approximately 4-5 days, mice became comfortable running in the darkness in the AVR.

Mice then began training on an active version of AVR1, where they needed to stop (defined as displacement less than 1.2 cm within 1 second) in the reward zone (the same zone in the passive AVR1 task) to retrieve water reward. Firstly, mice were stopped manually by experimenters for the first 2-3 days. After mice showed changes in behaviors (make frequent stops along the tracks), they were then left on their own. Experimenters provided continuous reinforcement learning throughout training sessions to help mice improve faster, including: (1) if mice missed too many rewards in a row, experimenters would start manually stopping the mice for ∼5 runs; (2) if mice slowed down in the correct reward zone but had trouble making a full stop, we switched them to an easier task that required stopping for only 0.5 second instead of 1 second, which usually was easy enough for the mice to do. Same as case (1), experimenters would sometimes provide manual stops if they thought it would be useful for mice’s improvement. Most mice were able to perform the active task in AVR1 after 2-3 weeks of training. Criteria used to determine their behaviors was the reward-retrieval-rate (>50% success rate for multiple consecutive sessions). Mice then started their imaging sessions.

#### Imaging AVR1 versus NA track

For each imaging day, mice started from no-auditory cue track (NA track, in darkness), in which no auditory cues were given that help mice in retrieving the reward, but they could still retrieve one if they correctly stopped in the reward zone. Mice finished all imaging sessions (2-3 FOVs) in NA track then were directly switched to AVR. Mice were left head-fixed till they finished all imaging sessions of the day (∼1 hour in total).

#### Behavioral training in VVR1

Mice were trained for active-task in the visual VR (210 cm track – VVR1). Similar training procedures were conducted as described above in the “*Behavioral training in AVR1*” section. Mice reached a similar level of performance (success rate in getting reward) as theirs in AVR after approximately 5-8 days.

#### Imaging AVR1 and VVR1

We imaged mice in AVR1 and VVR1 in two different ways:

1. The same FOV, both modalities: for each imaging day, mice randomly started either from AVR1 or VVR1 to account for the fact that mice were most engaged in the task at the beginning of the day when they were most thirsty. Mice finished the imaging session for one environment and were switched to the other. The same imaging FOV was imaged in the two sessions so that the same set of cells were compared directly.
2. All FOVs, single modality: for each imaging day, mice finished all imaging sessions for either auditory or visual environments. On the following day, mice would finish the same imaging sessions in the environment in the other sensory modality. The same FOV across these 2 days would be aligned for comparison.

#### Training and imaging in AVR2 and VVR2

We let mice explore another set of metrically-equivalent environments (AVR2 and VVR2; which included 4 cues (3 from the previous set and 1 novel cue) and a reward zone following the same cue in AVR1/VVR1. The orders of other cues on the tracks were permuted. The same training procedure used in AVR1/VVR1 tracks was followed until the mice could reach >50% success rate in getting reward for multiple consecutive days. For this pair of environments, we only imaged all FOVs for one sensory modality on each day and align FOVs across days for comparison.

#### Imaging in VAVR

Cues in AVR2 and VVR2 were combined to create a multisensory track (VAVR), in which auditory and visual cues were delivered at nonoverlapping locations. There were 2 reward zones in VAVR, each followed the same auditory or visual cues that mice learned in AVR2 or VVR2 before, other cue orders were permuted. The three mice, which still responded to auditory cues in VAVR, were trained in the track. For this track, mice were left on their own from beginning. Since all three mice were well trained in AVR2 and VVR2, they quickly adapted to VAVR and received reward at high success rates (> 50% success rate for both auditory and visual rewards). Therefore, the imaging data from days 1 to 3 were all used in the analysis.

### Behavioral analysis

#### Reward-retrieval-rate (success rate)

The percentage of rewards retrieved over the number of runs for each session.

#### Time stopping along track

The average amount of time spent at each spatial bin per run along the track was calculated for each session. We tested the difference in time spent between AVR1 and NA tracks at individual spatial bins. For each bin, we performed two-tailed t-test to identify whether the two conditions had different stopping time.

### Two-Photon Imaging

Imaging was performed using an Ultima 2Pplus microscope (Bruker) configured with the above VR setup. A tunable laser (Coherent, Chameleon Discovery NX) set to a 920nm excitation wavelength was used. Laser scanning was performed using a resonant-galvo scanner (Cambridge Technology, CRS8K). GCaMP6f fluorescence was isolated using a bandpass emission filter (525/25 nm) and detected using GaAsP photomultiplier tubes (Hamamatsu, H10770PB). A 16x water-immersion objective (Nikon, MRP07220) was used with ultrasound transmission gel (Sonigel, refractive index: 1.335985; Mettler Electronics, 1844) as the immersion media.

The anterior-posterior (AP) and the medial-lateral (ML) angles of the prism (i.e., the angle of the surface of the prism along to the AP or ML direction of the mouse) relative to the head-fixed position of the mouse were measured prior to the first imaging session. The head plate holder and rotatable objective angles were set to align the objective with the prism in the AP and ML direction, respectively, such that the objective was parallel to the prism surface. A black rubber tubing was wrapped around the objective and imaging window to prevent light leakage into the objective.

Microscope control and image acquisition were performed using Prairie View software (Bruker). Raw data were converted to images using the Bruker Image-Block Ripping Utility. Imaging was performed at 30 Hz with 512 × 512 resolution for a 500 × 500 FOV. Average beam power at the front of the objective was typically 90−115 mW. Imaging and behavior data were synchronized as described above.

### Image Processing

Imaging data was down-sampled by a factor of three by taking the average of each consecutive block of 3 frames and processed as previously described using custom MATLAB scripts^54^. Motion correction was performed using cross-correlation based, rigid motion correction. Identification of regions of interest (ROIs) with correlated fluorescence changes was performed using principal component analysis combined with independent component analysis (active cells)^54^. The fluorescence time course of individual ROIs was then extracted. The fractional change in fluorescence with respect to baseline (ΔF/F) was calculated as (F(t) – F0(t)) / F0(t)^25^. For each cell, significant calcium transients were identified using amplitude and duration thresholds, such that the false-positive rate of significant transient identification was 1%^24^. A final ΔF/F including only the significant calcium transients was used for all further analysis.

The mean ΔF/F for a cell was calculated as a function of position along the track in 5-cm bins. Data points when the mouse was moving below a stopping threshold, which was the same stopping criteria used in the active task (<1.2 cm/s), were excluded from this analysis. Moreover, since MEC activity is also modulated by acceleration^63^, data points when the mouse’s acceleration were above 1.1 cm/s^2^ were excluded from this analysis.

The acceleration threshold was determined as follows. First, we calculated average ΔF/F and acceleration for each spatial bin along track (here we used a spatial bin of 3 cm width). Second, we calculated correlation between ΔF/F and acceleration in a sliding window of 5 spatial bins. We plotted the correlation values with average accelerations of individual windows (Figure S8). This relationship saturated at value of acceleration = 1.1 cm/s^2^. Therefore, we chose this value to be the threshold for further analysis.

Some artifactual ROIs were occasionally caused by light leak from visual VR to the imaging window. Those ROIs normally exhibited extreme non-circularity (the ratio between the major and minor axes of the ellipse, which was used to approximate the shape of the ROI, was greater than 3) and were removed based on this feature from further analysis.

### Cell Alignment

All imaging sessions for a given mouse were aligned pairwise, as previously described^64^, to identify common cell pairs between each pair of sessions. Pairwise alignments were combined to generate the full set of possible cell alignments for all imaging days. These possible alignments were manually checked to determine the set of cells identified in all imaging sessions. The cells that overlapped in two FOVs were determined as common cells between the two FOVs, and the other cells were unique cells in each FOV.

### Data Analysis

*Inclusion/ Exclusion of mice used in each section*.

1. For AVR1-NA (Figures 2 and S1), 5 out of 7 mice that showed no difference in success rate followed a correct or error run was used.
2. For all metrically-equivalent tracks (Figures 3-5, S2-5, and S8), one mouse was excluded due to poor performance in AVR2 and VVR2.
3. For VAVR (Figures 6-8, S6-7), 3 mice that were still responding to auditory cues in VAVR when their imaging started were used.

#### Percentage of commonly active cells and random level

For each aligned pair of FOVs, percentage of common cells was calculated by the formula: nC / (n1 + n2 – nC); where nC was the number of common cells in the aligned FOV pair, n1 and n2 were the number of active cells in the two FOVs.

To calculate the random level of cell overlap: firstly, we manually identified all visible cells in the imaged FOV (n). For each pair of aligned imaging sessions: we had n1 and n2 as the number of active cells of the two sessions, respectively. (1) For session 1, we randomly allocated n1 cells into n available slots (the n visible cells). (2) Similarly, for session 2, we randomly allocated n2 cells into the same n slots. (3) We then calculated the percentage of slots that were randomly chosen in both times. We repeated these steps 1000 times to generate a random level of commonly active cells between the two sessions.

#### Active cell percentage (Figure 6E)

For each aligned pair of FOVs (for example: FOV A and FOV B), we extracted the number of common cells between FOVs. The percentage of cells in A that are active in A-B pair was calculated as (number of common cells) / (total number of active cells in A).

#### Percentage of cells with fields along the track

To calculate significant spatial firing fields (“in-fields”), which were regions of the track with significantly consistent activity, the mean ΔF/F was compared to shuffles of the original ΔF/F as described previously^54^. Each shuffle was calculated such that the original spatial position of each time point was preserved, but the ΔF/F was shuffled by bisecting the full ΔF/F time course at a random time point and swapping the order of the resulting halves. The bisecting point was localized between the 5% and 95% of the dataset to ensure that the bisected activity was significantly different from the original activity^65^. The mean ΔF/F of the shuffle was then calculated as described above. An “in-field” was defined as a region of at least 3 consecutive 5cm bins (except that the fields at the beginning and end of the track could have 2 bins) that had a mean ΔF/F higher than 80% of 1000 shuffles at the corresponding bins. Additionally, at least 20% of runs were to have a significant calcium transient in the “in-field” region. For the activity of each cell on one day, a field vector (42, 66, and 140 elements corresponding to the 42, 66, 170 bins on 210-cm, 330-cm, and 700-cm tracks, respectively) was generated by using ones and zeros to indicate whether individual spatial bins were in-field or not in-field bins, respectively. The percentage of cells with fields was then calculated on a bin-by-bin basis by dividing the number of ones in each bin by the total number of cells.

#### Shuffled field distribution

For the field vector of each cell, we generated 1000 shuffles: in each shuffle, the vector was randomly bisected, and the resulting halves were swapped. Shuffled fields distribution was the average values within individual spatial bins across all shuffled data.

#### Difference in the percentage of cells with fields along the track

The difference in the percentage of cells with fields along track for two conditions (e.g., AVR1 and NA) was calculated for each spatial bin, where we performed two-tailed t-test to identify whether the two conditions had statistically different number of cells with fields in the bin.

#### Activity matrix

For each cell, a spatially binned mean ΔF/F for each cell was averaged across all runs along the track, generating a 1D array (ΔF/F vector). In-field-amplitude was calculated as follows: for each cell’s ΔF/F vector, the values within spatial field areas was kept as original values, whereas those outside of field areas were set to zero. This calculation generated the field-only ΔF/F, which was further normalized between 0 and 1. The normalized field-only ΔF/F vectors of individual cells were then sorted based on their peak locations and ordered in the two-dimensional (2D) matrix (size = total number of cells × number of spatial bins), representing the in-field-amplitude distributions of all cells.

#### Nearest neighbor (NN) distance analysis

To calculate anatomical distances between two cell groups A and B, we first determined the group with less cells (e.g., group A), and randomly subsampled the other group (group B) to be sample-matched with A. For a given number of NNs (e.g., 5 neighbors), we calculated the distances between individual cells in group A with their nearest 5 cells in subsampled group B (AB distances). The distances of cells in subsampled group B to group A cells (BA distances) were similarly calculated. The distance between groups A and this subsampled group B was the mean of AB and BA distances. The above random subsampling of group B was conducted 100 times and the final distance between groups A and B under this particular number of NNs was the mean of all the 100 distances.

To calculate the shuffled distance for each set of group A with subsampled group B, cells were randomly permuted among group A and subsampled group B for 100 times, and the distance between the two shuffled groups was similarly calculated as described above.

The above procedure was repeated for different numbers of NNs. For the cells active in AVR versus in VVR, the minimal number of NNs was 5. For auditory cells versus visual cells and auditory reward cells versus visual reward cells in VAVR, the minimal number of NNs was 1, because there were fewer cells in each group. The maximal number of NNs for a pair of groups A and B was the number of cells in A, given that group A had less cells than group B.

To determine whether groups A and B were anatomically separated, their distances in all FOVs and under all NNs were pooled and compared with shuffles, as represented by the cumulative density plot in Figures 3J. If groups A and B were anatomically separated, their distances should be longer than shuffles. If the cells were anatomically intermingled, their distances should be comparable to shuffles.

#### Ratio of the percentage of fields in cue near and away from reward

For each FOV, field vector of all cells was grouped in a 2D matrix (size = number of cells × number of spatial bins). The matrix columns that corresponded to each cue area was used to calculate the average percentage of cells with fields in each cue area by taking average of all value of those matrix columns. Ratio between near and far away from reward was calculated using the values associated with cues that were near and away from the reward.

#### Run-by-Run (RBR) Activity Consistency

The run-by-run consistency for a given cell in a particular imaging session was calculated as previously described^66^. The spatially binned mean ΔF/F for each run along the track was correlated with that of every other run. The average of these correlations was the run-by-run consistency value for a given cell.

#### RBR consistency at cues (Cue consistency)

The RBR consistency within a cue area (auditory or visual zone) was calculated in the same manner described above except using only ΔF/F of the track that was associated with the cue area.

#### RBR consistency difference between Correct vs Error runs

Due to the high success rate, there usually was a smaller number of error runs in an imaged session, we calculated RBR consistency for correct versus error runs as follows:

1. If there was no more than 1 error run, we only calculated RBR consistency in correct runs, and error run consistency were set to Nan.
2. If there were more than 6 error runs, which was the smallest number of correct runs among all sessions: we divided the session into correct and error runs and separately calculated their RBR consistency.
3. If there were less than 6 but more than 1 error runs: in this case, a session normally had a large number of correct runs and a very small number of error runs. To account for potential RBR consistency difference caused by the large difference in the numbers of correct and error runs, we performed the following steps: (a) we used all ΔF/F of the error runs (e.g., n runs) to calculate run consistency as described above. (b) For correct runs, we used ΔF/F of a subset of n consecutive correct runs, starting fr^om^ 1st correct run (1 to n, 2 to (n+1), …) to calculate RBR consistency of the subset, and further averaged the consistency of all subsets for the final RBR consistency of correct runs.

#### Normalized cue consistency difference between the areas near and away from reward

For each cell: (1) we calculated the cue consistency for each cue area; (2) these values were then normalized to range 0−1 to account for difference in RBR consistency between AVRs and VVRs; (3) we averaged the values for all cues near or away-from reward. (4) Finally, we subtracted the value of near-reward from the value of away-from reward. The result was the average of normalized difference across all cells.

#### Activity correlation

For commonly active cells in any metrically equivalent pair (AVR1-VVR1; AVR2-VVR2), we had their mean ΔF/F as a function of track position in both environments. For each cell, we calculated correlation between their mean ΔF/F in each environment. Distribution of activity correlation was performed by grouping all values across all cells. All correlations were compared with randomly shuffled distribution created by permuting activity patterns between cells. The comparison of distributions of real data and shuffles was made by two-tailed Kolmogorov-Smirnov test.

#### Cue score and cue cell classification

Cue scores were calculated as previously described^10^. Due to the short track, we only allowed the cell activity to be shifted up to 4 spatial bins (20 cm) relatively to the “cue template (type A, for cue score calculation)”, which was a linear vector consisted of zeros and ones that represented spatial bins outside and within cue areas (6 bins per cue), respectively. Cue score calculation on metrically-equivalent tracks used the identical cue template and spatial shift range. Note that cue template A was specific for cue score calculation and was different from the “cue template B” for cue correlation below.

Cue score thresholds were calculated as the 95^th^ percentile of 1000 shuffles of each cell from all mice and imaging sessions in a given environment. The shuffles were performed as described in “*Percentage of cells with fields along the track”.* If the cue score of a cell was above the cue score threshold, the cell was classified as a cue cell.

#### Cue cell percentage and shuffled shared cue cell percentage

For each aligned pair of FOV, we identified cue cell as described above for each FOV. Shared cue cell percentage is calculated as nCC / (n1 + n2 – nCC). nCC was the number of cells that were identified as cue cell in both FOVs, n1 and n2 were the numbers of active cells in each FOV.

Shuffled shared cue cell percentage was calculated by: (1) permuting the order of cells so that cue cells of the same cell on a pair of FOVs became misaligned; (2) shuffled cue cell percentage was calculated as nSCC / (n1 + n2 – nSCC); nSCC was the number of cells that were identified as cue cell in both FOVs after being misaligned, n1 and n2 were the numbers of active cells in each FOV, identical to those mentioned above. We repeated this 1000 times so that a distribution of shared cue cell percentage at random level was created.

To determine whether two tracks shared a significant percentage of cue cells, we calculated p_above, which was the percentage of “shuffle percentage” data that were above the “real percentage”. For example, p_above = 0.045 means that 45 out of 1000 shuffle percentages were above the real percentage. If p_above was not above 0.05, we consider that the two tracks had a significant population of shared cue cells.

#### Cue Correlation

Cue correlation was the correlation between mean ΔF/F of each cell along the track and the cue template (type B, for cue correlation calculation), which was a linear vector consisted of zeros and ones that represented spatial bins outside and within cue areas, respectively. Since visual cue cells could have spatially shifted response relative to cues^10^, we expanded the cue areas in cue template B to 10 spatial bins (instead of 6 bins in cue template A for cue score calculation above) to account for the shift.

#### Sensory score calculation in VAVR track

Sensory score was calculated using mean ΔF/F of each cell along the track. Auditory cue zone contained 40 spatial bins, which consisted of 10 bins centered around each auditory cue and there was a total of 4 cues. Similarly, visual cue zone also contained 40 bins for the 4 visual cues. For each cell, mean ΔF/F values in auditory (A) or visual cue zones (V) were first calculated and a sensory score was defined as the difference between A and V normalized by the sum of A and V.

Shuffled sensory scores were calculated by randomly permuting mean ΔF/F of each cell for 100 times and conducting the above calculation on the permuted ΔF/F.

Fake sensory scores were calculated by first randomly assigning the 8 cues into “fake” auditory (AF) and visual (VF) cue groups, and then calculating sensory score as the difference between AF and VF normalized by the sum of AF and VF.

#### Distribution of sensory scores

Since sensory scores were distributed from −1 and 1, this range was divided into 21 bins and the distribution of sensory scores (real scores, shuffled scores, and scores with other cue combinations) was calculated as the number of scores/cells distributed in each bin.

#### Identification of auditory and visual cells in VAVR track

Auditory and visual cells were identified as the cells with sensory scores above 99^th^ and below 1^st^ percentiles of shuffled sensory scores mentioned above, respectively.

#### Reward score calculation in unisensory and VAVR tracks

Similar to the calculation of sensory scores, mean ΔF/F values in reward (R) or no-reward zones (N) were first calculated, and a reward score was defined as the difference between R and N normalized by the sum of R and N. In VAVR track, the two reward zones were combined.

Shuffled reward scores were calculated by randomly permuting mean ΔF/F of each cell for 100 times and conducting the above calculation on the permuted ΔF/F.

#### Identification of reward cells on unisensory and VAVR tracks

Reward cells were identified as the cells with reward scores above 99^th^ percentiles of shuffled reward scores mentioned above. Common reward cells across two unisensory tracks (e.g., AVR1 and VVR1) were the cells that passed the thresholds on both tracks.

To determine whether two unisensory tracks shared a significant population of reward cells (Figure S5), the percentage of common reward cells among the total number of common cells in the two tracks was first calculated (real percentage). To calculate the shuffle percentage, reward scores of common cells were randomly permuted, and a shuffled percentage of common reward cells was recalculated (shuffle percentage). The shuffles were conducted 1000 times and the p_above was the percentage of “shuffle percentage” data that were above the “real percentage”. For example, p_above = 0.045 means that 45 out of 1000 shuffle percentages were above the real percentage. If p_above was not above 0.05, we consider that the two tracks had a significant population of common reward cells.

#### Sensory reward score calculation on VAVR track

For each reward cell, mean ΔF/F values in auditory reward (A) or visual reward zones (V) were first calculated, and a sensory reward score was defined as the difference between A and V normalized by the sum of A and V.

#### Identification of visual and auditory reward cells on VAVR track

Sensory reward scores of real cells were compared with those of shuffles, which were constructed by randomizing the activity of reward cells in the auditory and visual reward zones. There were 1000 shuffles made for each cell so that a distribution of shuffled sensory reward scores was created. The reward cells with sensory reward scores above the 99^th^ percentile and below the 1^st^ percentile of the shuffled distribution were identified as auditory and visual reward cells, respectively.

#### General statistics

Linear correlations and the corresponding r and p values were calculated using a two-tailed Pearson’s linear correlation coefficient. The comparison of spatial field distribution and time in stops along the track was done by two-tailed t-tests for both tails to identify which distribution was different at each specific spatial bin. Other significant values were calculated using Student’s paired t-test (i.e., for commonly active population) or two-sample t-test (i.e., for uniquely active population). p values less than or equal to 0.05 were considered significant (* ≤ 0.05, ** ≤ 0.01, *** ≤ 0.001). All figures show mean ± SEM, except otherwise noted.

---

**Figure S1:**
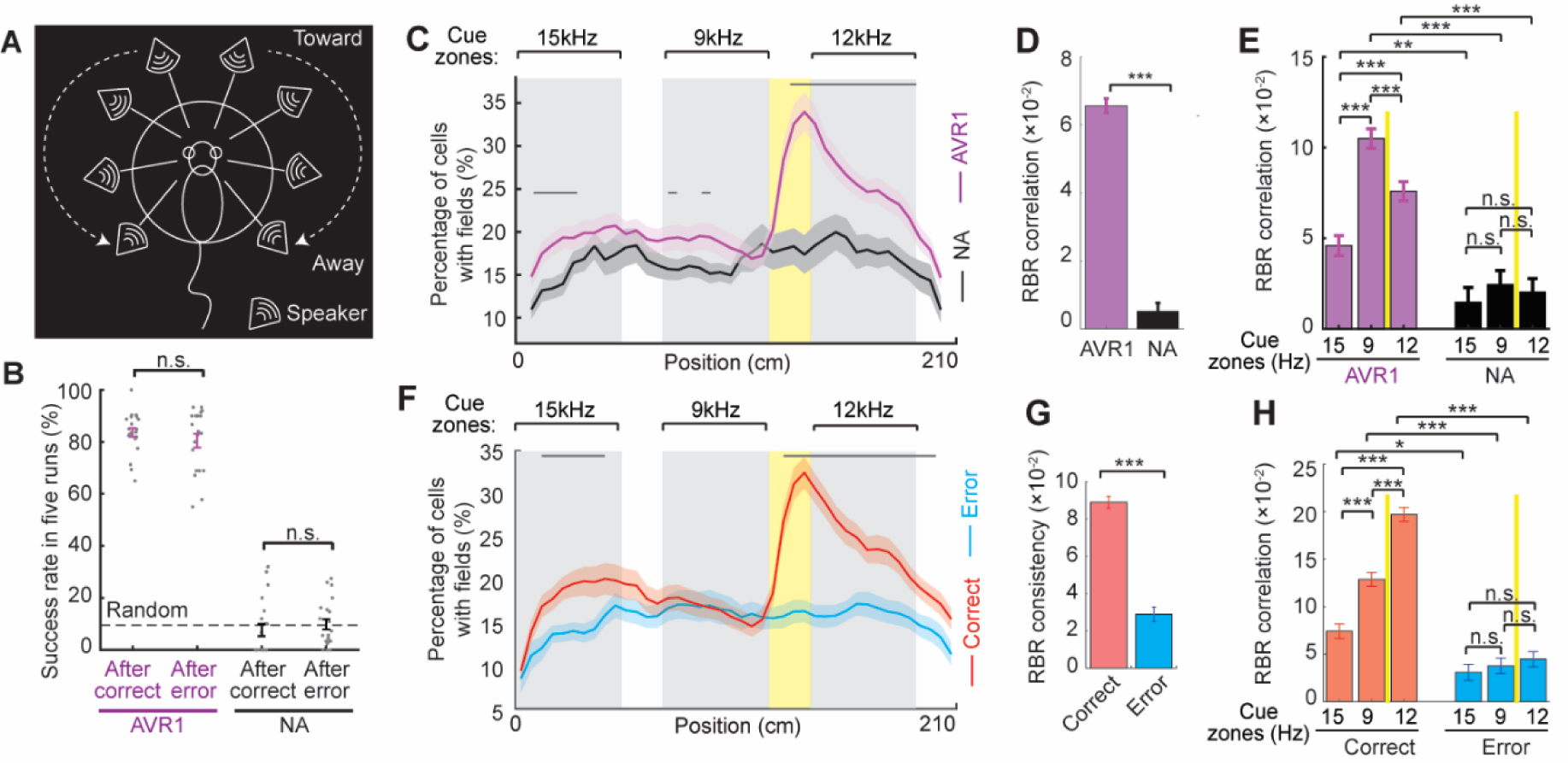
AVR setup, behavior and MEC response in AVR1 and NA A. Eight speakers positioned in a circle with the mouse head in the center. Auditory cues were played from the front to the back speakers to mimic sound orientation effect when mice ran toward and away from a sound cue. B. Success rate of receiving the reward within 5 runs after a correct or an error run in AVR1 and NA. The dashed line indicates the random level of success rate. C. Percentage of unique cells in AVR1 (magenta) or NA (black) with spatial fields at each spatial bin along the track. D. RBR activity consistency of unique cells in AVR1 or NA. p value = 1.2×10^−65^. E. RBR activity consistency of unique cells in AVR1 or NA at individual auditory cues. p values for the difference between AVR1 and NA in 15 kHz cue: 0.0014; 9 kHz cue: 7.7×10^−18^, and 12 kHz cue: 1.1×10^−9^. p values between cues within AVR1: 15 kHz versus 9 kHz: 1.7×10^−14^; 15 kHz versus 12 kHz: 7.9×10^−5^, and 9 kHz versus 12 kHz: 1.1×10^−4^. F. Percentage of AVR1 unique cells with spatial fields along the track in correct (red) and error runs (blue). G. RBR activity consistency of AVR1 unique cells in correct and error runs. p =6.9×10^−26^. H. RBR activity consistency of AVR1 unique cells in each auditory cue zones during correct and error runs. p values for the difference between correct and error runs at 15 kHz cue: 0.0125; 9 kHz cue: 3.4×10^−10^, and 12 kHz cue: 2.6×10^−23^. p values between cues within correct runs: 15 kHz versus 9 kHz: 1.0×10^−5^; 15 kHz versus 12 kHz: 1.2×10^−21^, and 9 kHz versus 12 kHz: 2.6×10^−9^. *p ≤ 0.05, **p ≤ 0.01, ***p ≤0.001, n.s. p > 0.05. Error bars represent mean ± SEM. Data was collected from 5 mice.

**Figure S2:**
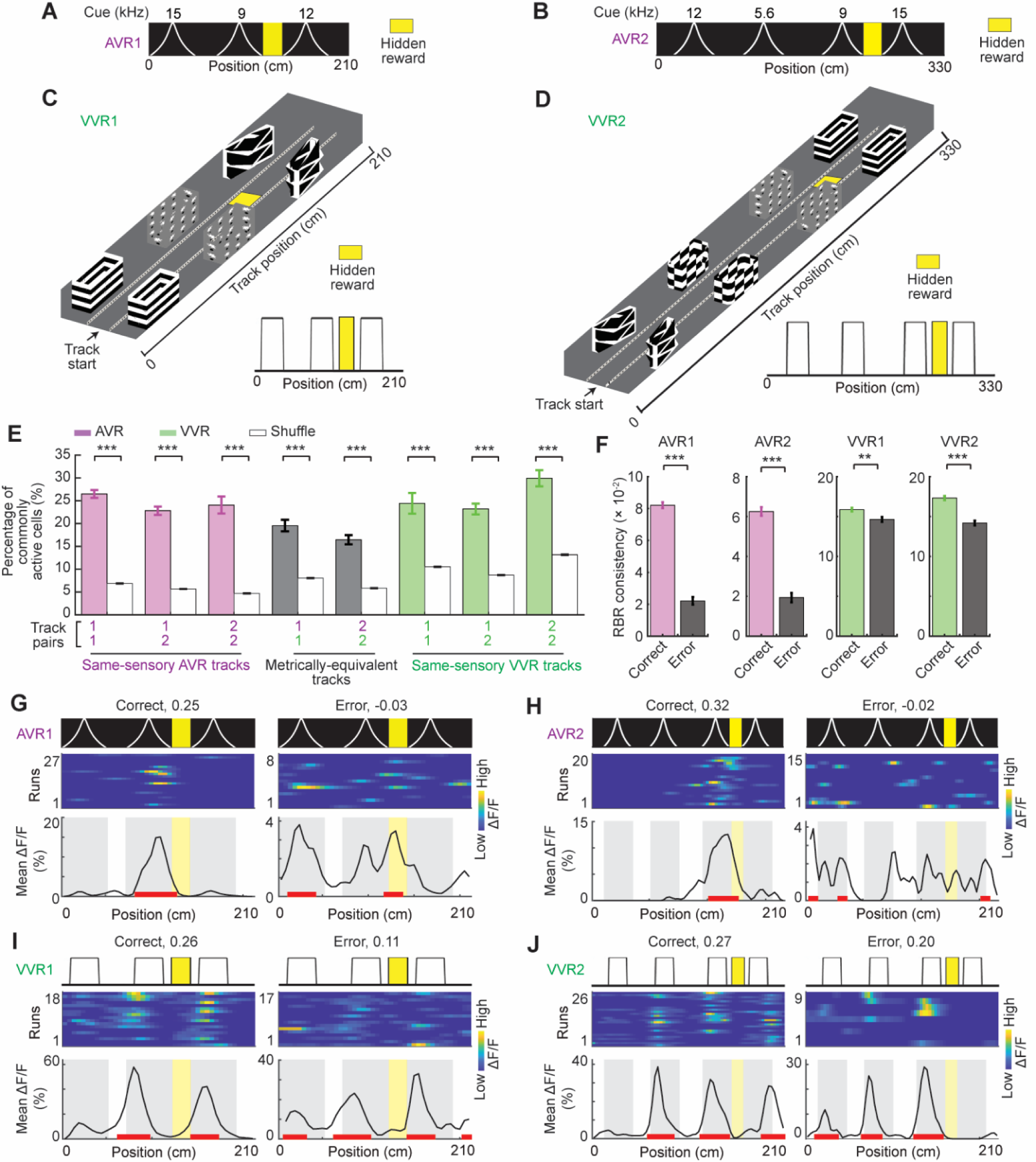
Metrically-equivalent track arrangement and cell activity features. A. AVR1 track. B. AVR2 track. Three cues in AVR2 were identical to the ones on AVR1. AVR2 also had a new cue (5.6 kHz). C. VVR1 track and its simplified diagram. Yellow indicates a hidden reward zone. D. VVR2 track and its simplified diagram. Three cues in VVR2 were identical to the ones on VVR1. VVR2 also had a new cue. Yellow indicates a hidden reward zone. E. Percentage of commonly active cells between pairs of tracks, in comparison to shuffles. p values (from left to right): 3.6×10^−43^; 6.9×10^−38^; 4.2×10^−11^; 7.5×10^−10^; 3.7×10^−15^; 3.5×10^−5^; 1.8×10^−17^; and 2.8×10^−10^. F. RBR activity consistency between correct and error runs in all four tracks. p values for AVR1: 8.5×10^−71^; AVR2: 2.6×10^−26^; VVR1: 0.0023; VVR2: 2.4×10^−10^. G-J. Examples for RBR activity consistencies of the same cell in correct and error runs (the consistency values are on top) in the four tracks in A-D. *p ≤ 0.05, **p ≤ 0.01, ***p ≤0.001, n.s. p > 0.05. Error bars represent mean ± SEM. Data was collected from 6 mice.

**Figure S3:**
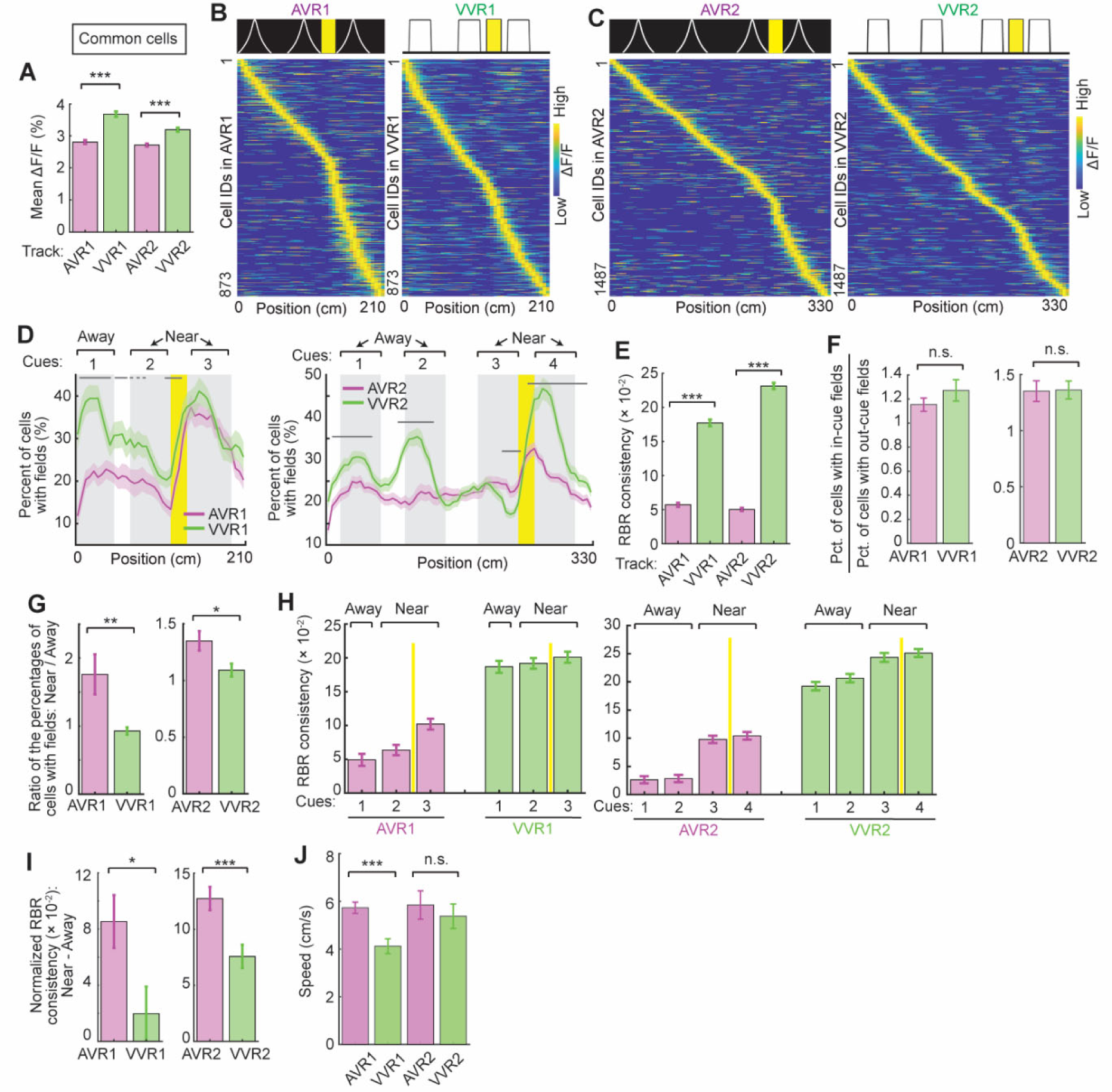
Different response in common cells in metrically-equivalent tracks A. Average calcium response (mean ΔF/F) of common cells in the two pairs of metrically-equivalent tracks. p values: AVR1 versus VVR1: 6.9×10^−17^; AVR2 versus VVR2: 5.5×10^−10^. B. Activity matrices of common cells in AVR1 (left) and VVR1 (right). C. Activity matrices of common cells in AVR2 (left) and VVR2 (right). D. Percentage of common cells in AVR1 and VVR1 (left) and in AVR2 and VVR2 (right) with spatial fields along the track. E. RBR activity consistency of common cells in metrically-equivalent tracks. p values for AVR1 versus VVR1: 6.9×10^−89^; for AVR2 versus VVR2: 7.3×10^−226^. F. Ratio of the percentages of common cells with in-cue and out-cue fields. G. Ratio of the percentages of common cells with fields at cues that are near and away from reward. Left: AVR1 versus VVR1, p = 0.0072; right: AVR2 and VVR2, p = 0.0133. H. RBR activity consistency in cue areas for common cells in AVR1 and VVR1 (left) and in AVR2 and VVR2 (right). I. Normalized difference in RBR activity consistency at cues near and away from reward for common cells in AVR1 and VVR1 (p = 0.0144) and in AVR2 and VVR2 (p = 5.7×10^−4^). J. Speed in all four tracks. p value for AVR1 and VVR1: 4.1×10^−4^. The speed was calculated only in correct runs. *p ≤ 0.05, **p ≤ 0.01, ***p ≤0.001, n.s. p > 0.05. Error bars represent mean ± SEM. Data was collected from 6 mice.

**Figure S4:**
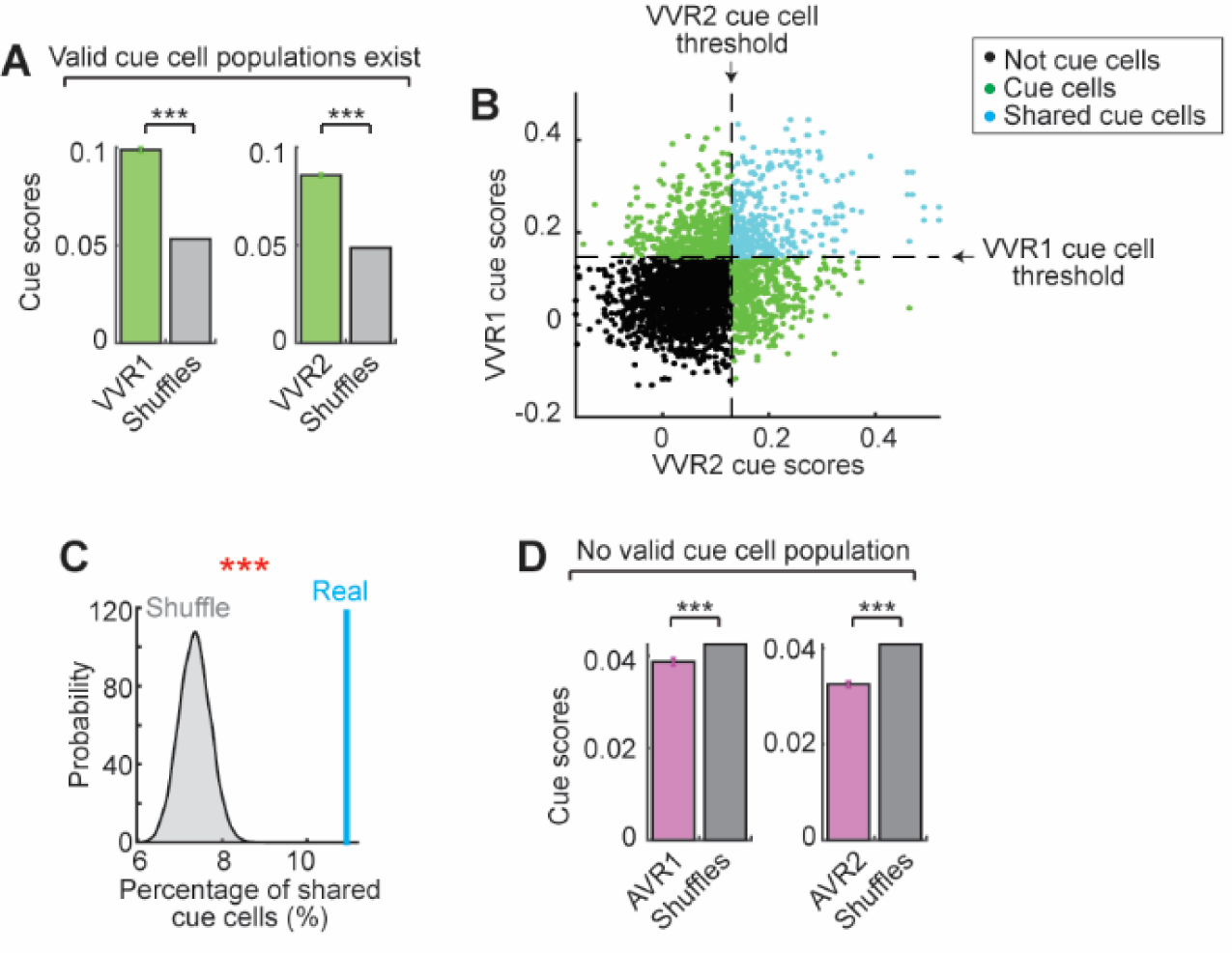
Cue cells were identified in VVRs, but not in AVRs A. Comparison of cue scores for common cells in VVR1 and VVR2 with shuffles. p values for VVR1: 0; for VVR2: 0. The significantly higher scores of real cells suggest the existence of specific cue cells in VVRs. B. Common cells in VVR1 and VVR2 identified as cue cells (green) and shared cue cells in the two tracks (cyan). C. The percentage of shared cue cells between VVR1 and VVR2 (real, cyan line) is higher than shuffle percentages (gray). To calculate the shuffle percentage, cue scores of common were randomly permuted, and a shuffled percentage of shared cue cells was recalculated (shuffle percentage). The shuffles were conducted 1000 times and the p_above was the percentage of “shuffle percentage” data that were above the “real percentage”. For example, p_above = 0.04 means that 40 out of 1000 shuffle percentages were above the real percentage. If p_above was not above 0.05, we consider that the two tracks had a significant population of shared cue cells. In this case, p_above = 0. D. Comparison of cue scores for common cells in AVR1 and AVR2 with shuffles. p values for AVR1: 4.7×10^−8^; for AVR2: 5.1×10^−62^. The significantly lower scores in real cells suggest the absence of cue cells in AVRs. *p/p_above ≤ 0.05, **p/p_above ≤ 0.01, ***p/p_above ≤0.001, n.s. p/p_above > 0.05. Error bars represent mean ± SEM. Data was collected from 6 mice.

**Figure S5.**
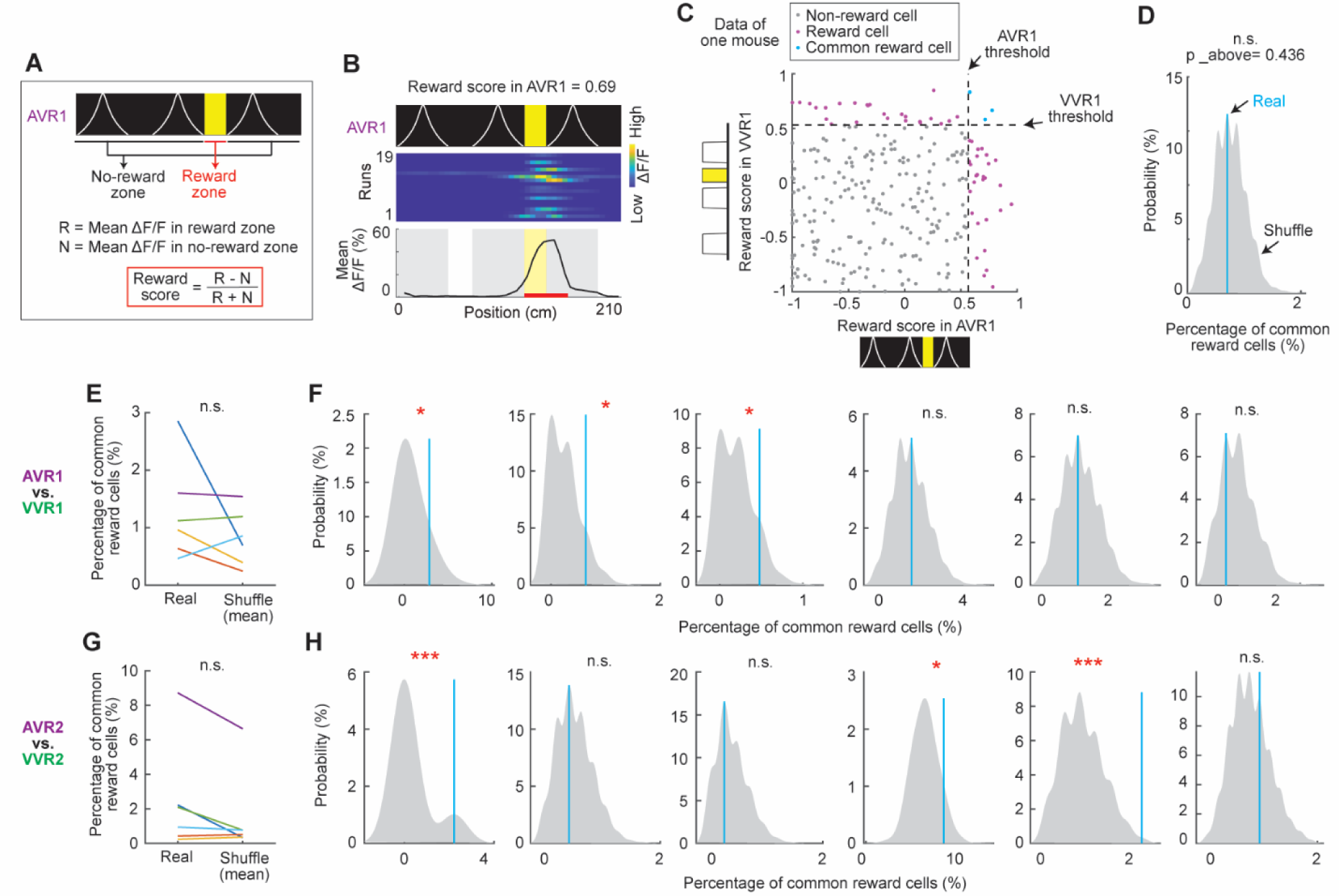
Reward encoding in metrically-equivalent unisensory tracks A. Calculation of reward score. B. An example cell with high reward score in AVR1. C. Classification of common reward cells in AVR1 and VVR1. The common reward cells were the ones passing reward score thresholds (99^th^ percentile of shuffles) in both tracks. D. The percentage of common reward cells in AVR1 and AVR2 (blue) compared to shuffle percentages (gray). The shuffle percentages were generated as described in Figure S4C. If p_above was not higher than 0.05, the two tracks had a significant population of common reward cells. In this case, p_above = 0.436. E. The comparison of the percentage of common reward cells of AVR1 and VVR1 in real data (real) with the mean of shuffle percentages calculated in D (Shuffle (mean)). Each line represents one mouse. F. Similar to D but the comparison was made using the data of AVR1 and VVR1 of the individual 6 mice. Significant p_above values from left to right: 0.018, 0.039, 0.032. G. Similar to E but for AVR2 and VVR2. H. Similar to F but for AVR2 and VVR2. Significant p_above values from left to right: 0, 0.05, 0.004. *p_above ≤ 0.05, **p_above ≤ 0.01, ***p_above ≤0.001, n.s. p_above > 0.05. Data was collected from 6 mice.

**Figure S6:**
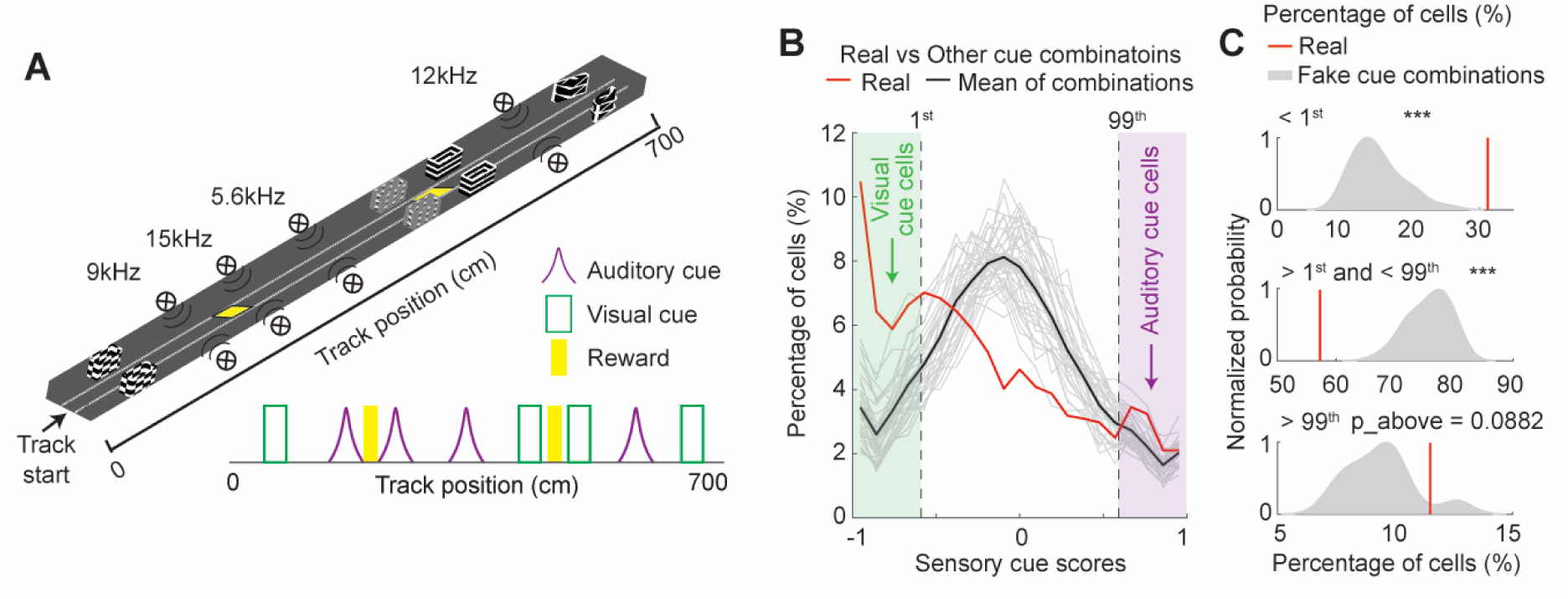
MEC neural response in VAVR A. VAVR linear track and its simplified diagram. B. Similar to Figure 6H but for the distributions of real sensory cue scores and fake scores using random cue combinations (34 combinations: gray; mean: black). C. Comparisons between real (red) and fake cases (gray) in the percentages of cells with sensory cue scores below 1^st^ threshold (p_above = 0), between 1^st^ and 99^th^ thresholds (p_below = 0), and above 99^th^ threshold (p_above = 0.0882). *p ≤ 0.05, **p ≤ 0.01, ***p ≤0.001, n.s. p > 0.05. Error bars represent mean ± SEM. Data was collected from 3 mice.

**Figure S7.**
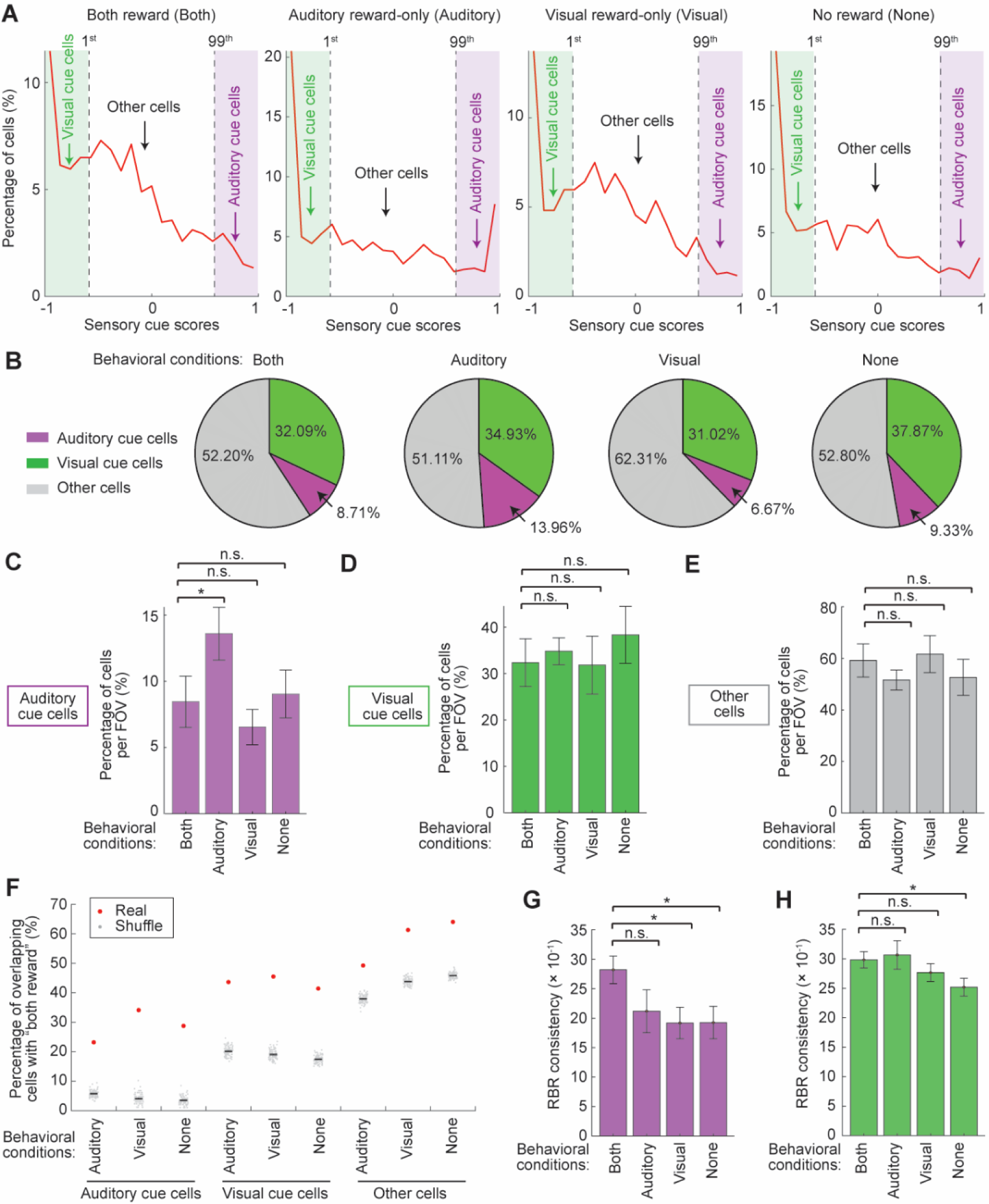
Visual and auditory cue cells in different behavioral conditions in VAVR A. Distributions of sensory cue scores (red curve) in four behavioral conditions (from left to right), including the runs when mice received both rewards (Both, similar to Figure 6H), only auditory reward (Auditory), only visual reward (Visual), and no reward (None). For each plot, vertical dashed lines indicate the 1^st^ and 99^th^ percentiles of shuffle scores in Figure 6H, which were used to identify visual (in green zone) and auditory cue cells (in magenta zone), respectively. The cells between the dashed lines are “other” cells. B. Percentages of other cells (gray), visual cue cells (green), and auditory cue cells (magenta) in data from the four behavioral conditions in A. C-E. Percentages of auditory cue cells (C), visual cue cells (D), and other cells (E) in data from the above four conditions calculated per FOV. For (C), p value for “Both” versus “Auditory”: 0.0469. F. Percentages of overlapping cells between “Both” and the other three conditions (Auditory, Visual, and None, red dots) for the three cell categories (auditory, visual, and other cells). These percentages were compared with 100 sets of shuffles (gray dots), which were the percentages of overlapped cells after randomly selecting auditory, visual, and other cells according to the cell numbers in the real data for all four conditions. Black error bars indicate mean and standard errors of the shuffles. In all cases, the real percentage was much higher than those of shuffles, indicating that in real case, the three cell categories across different run conditions significantly overlapped. G. RBR activity consistency of auditory cue cells in the four behavioral conditions. p values for “Both” versus “Visual”: 0.0122; for “Both” versus “None”: 0.0153. H. RBR activity consistency of visual cue cells in the four behavioral conditions. p values for “Both” versus “None”: 0.0244. *p ≤ 0.05, **p ≤ 0.01, ***p ≤0.001, n.s. p > 0.05. Error bars represent mean ± SEM. Data was collected from 3 mice.

**Figure S8:**
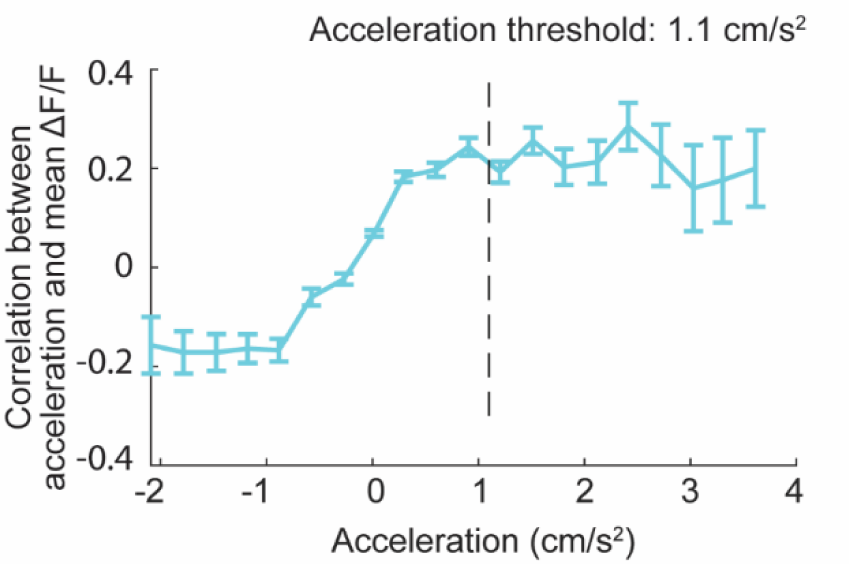
Correlation between acceleration and mean ΔF/F. Data was binned by dividing the range from smallest acceleration value to largest acceleration value equally into 20 bins. Error bars indicate mean ± SEM for correlation values within individual bins. The vertical dashed line indicates the acceleration threshold used in data analysis. Data was collected from the 6 mice in metrically-equivalent tracks.

**Table S1.** P values for the comparison between the percentage of commonly active cells in Figure 3G. As an example, AVR1-AVR1 indicates the percentage of cells commonly active in two imaging sessions in AVR1 (a), AVR1-VVR1 indicates the percentage of cells commonly active in between on session in AVR1 and another session in VVR1 (b). AVR1-AVR1 vs. AVR1-VVR1 means the p values for the comparison of a and b. Data was collected from 6 mice.

| Pair | p value |
| --- | --- |
| AVR1-AVR1 vs. AVR1-VVR1 | $7.4 \times 10^{-6}$ |
| AVR1-AVR1 vs. AVR2-VVR2 | $4.7 \times 10^{-12}$ |
| AVR1-AVR2 vs. AVR1-VVR1 | 0.0124 |
| AVR1-AVR2 vs. AVR2-VVR2 | $6.9 \times 10^{-6}$ |
| AVR2-AVR2 vs. AVR1-VVR1 | 0.0219 |
| AVR2-AVR2 vs. AVR2-VVR2 | $7.4 \times 10^{-4}$ |
| VVR1-VVR1 vs. AVR1-VVR1 | 0.0376 |
| VVR1-VVR1 vs. AVR2-VVR2 | 0.0046 |
| VVR1-VVR2 vs. AVR1-VVR1 | 0.0129 |
| VVR1-VVR2 vs. AVR2-VVR2 | $3.3 \times 10^{-5}$ |
| VVR2-VVR2 vs. AVR1-VVR1 | $5.3 \times 10^{-6}$ |
| VVR2-VVR2 vs. AVR2-VVR2 | $3.5 \times 10^{-8}$ |

## References

1. Bruns, P., and Roder, B. (2023). Development and experience-dependence of multisensory spatial processing. Trends Cogn Sci. 10.1016/j.tics.2023.04.012.

2. Bates, S.L., and Wolbers, T. (2014). How cognitive aging affects multisensory integration of navigational cues. Neurobiol Aging 35, 2761–2769. 10.1016/j.neurobiolaging.2014.04.003.

3. Epstein, R.A., Patai, E.Z., Julian, J.B., and Spiers, H.J. (2017). The cognitive map in humans: spatial navigation and beyond. Nat Neurosci 20, 1504–1513. 10.1038/nn.4656.

4. O’Keefe. J.; Nadel, L. (1978). The hippocampus as a cogintive map. Oxford: Clarendon Press.

5. Leinonen, H., and Tanila, H. (2018). Vision in laboratory rodents-Tools to measure it and implications for behavioral research. Behav Brain Res 352, 172–182. 10.1016/j.bbr.2017.07.040.

6. Hoy, J.L., Yavorska, I., Wehr, M., and Niell, C.M. (2016). Vision Drives Accurate Approach Behavior during Prey Capture in Laboratory Mice. Curr Biol 26, 3046–3052. 10.1016/j.cub.2016.09.009.

7. Koay, S.A., Thiberge, S., Brody, C.D., and Tank, D.W. (2020). Amplitude modulations of cortical sensory responses in pulsatile evidence accumulation. Elife 9. 10.7554/eLife.60628.

8. Terry, A.V., Jr. (2009). Spatial Navigation (Water Maze) Tasks. In Methods of Behavior Analysis in Neuroscience, J.J. Buccafusco, ed.

9. Hafting, T., Fyhn, M., Molden, S., Moser, M.B., and Moser, E.I. (2005). Microstructure of a spatial map in the entorhinal cortex. Nature 436, 801–806. 10.1038/nature03721.

10. Kinkhabwala, A.A., Gu, Y., Aronov, D., and Tank, D.W. (2020). Visual cue-related activity of cells in the medial entorhinal cortex during navigation in virtual reality. Elife 9. 10.7554/eLife.43140.

11. Tukker, J.J., Beed, P., Brecht, M., Kempter, R., Moser, E.I., and Schmitz, D. (2022). Microcircuits for spatial coding in the medial entorhinal cortex. Physiol Rev 102, 653–688. 10.1152/physrev.00042.2020.

12. Heffner, H.E., and Heffner, R.S. (1985). Hearing in two cricetid rodents: wood rat (Neotoma floridana) and grasshopper mouse (Onychomys leucogaster). J Comp Psychol 99, 275–288.

13. Turner, J.G., Parrish, J.L., Hughes, L.F., Toth, L.A., and Caspary, D.M. (2005). Hearing in laboratory animals: strain differences and nonauditory effects of noise. Comp Med 55, 12–23.

14. Zheng, Q.Y., Johnson, K.R., and Erway, L.C. (1999). Assessment of hearing in 80 inbred strains of mice by ABR threshold analyses. Hear Res 130, 94–107. 10.1016/s0378-5955(99)00003-9.

15. Rossier, J., Haeberli, C., and Schenk, F. (2000). Auditory cues support place navigation in rats when associated with a visual cue. Behav Brain Res 117, 209–214. 10.1016/s0166-4328(00)00293-x.

16. Watanabe, S., and Yoshida, M. (2007). Auditory cued spatial learning in mice. Physiol Behav 92, 906–910. 10.1016/j.physbeh.2007.06.019.

17. Funamizu, A., Kuhn, B., and Doya, K. (2016). Neural substrate of dynamic Bayesian inference in the cerebral cortex. Nat Neurosci 19, 1682–1689. 10.1038/nn.4390.

18. Gao, S., Webb, J., Mridha, Z., Banta, A., Kemere, C., and McGinley, M. (2020). Novel Virtual Reality System for Auditory Tasks in Head-fixed Mice. Annu Int Conf IEEE Eng Med Biol Soc 2020, 2925–2928. 10.1109/EMBC44109.2020.9176536.

19. Vinogradov, O.S. (1975). Functional orgnization of the limbic system in the process of registration of information: facts and hypothesis. The hippocampus.

20. Aronov, D., Nevers, R., and Tank, D.W. (2017). Mapping of a non-spatial dimension by the hippocampal-entorhinal circuit. Nature 543, 719–722. 10.1038/nature21692.

21. Derner, M., Chaieb, L., Dehnen, G., Reber, T.P., Borger, V., Surges, R., Staresina, B.P., Mormann, F., and Fell, J. (2021). Auditory Beat Stimulation Modulates Memory-Related Single-Neuron Activity in the Human Medial Temporal Lobe. Brain Sci 11. 10.3390/brainsci11030364.

22. Xiao, C., Liu, Y., Xu, J., Gan, X., and Xiao, Z. (2018). Septal and Hippocampal Neurons Contribute to Auditory Relay and Fear Conditioning. Front Cell Neurosci 12, 102. 10.3389/fncel.2018.00102.

23. Wan, H., Warburton, E.C., Kusmierek, P., Aggleton, J.P., Kowalska, D.M., and Brown, M.W. (2001). Fos imaging reveals differential neuronal activation of areas of rat temporal cortex by novel and familiar sounds. Eur J Neurosci 14, 118–124. 10.1046/j.0953-816x.2001.01625.x.

24. Heys, J.G., Rangarajan, K.V., and Dombeck, D.A. (2014). The functional micro-organization of grid cells revealed by cellular-resolution imaging. Neuron 84, 1079–1090. 10.1016/j.neuron.2014.10.048.

25. Low, R.J., Gu, Y., and Tank, D.W. (2014). Cellular resolution optical access to brain regions in fissures: imaging medial prefrontal cortex and grid cells in entorhinal cortex. Proc Natl Acad Sci U S A 111, 18739–18744. 10.1073/pnas.1421753111.

26. Wilming, N., Konig, P., Konig, S., and Buffalo, E.A. (2018). Entorhinal cortex receptive fields are modulated by spatial attention, even without movement. Elife 7. 10.7554/eLife.31745.

27. Meister, M.L.R., and Buffalo, E.A. (2018). Neurons in Primate Entorhinal Cortex Represent Gaze Position in Multiple Spatial Reference Frames. J Neurosci 38, 2430–2441. 10.1523/JNEUROSCI.2432-17.2018.

28. Killian, N.J., Jutras, M.J., and Buffalo, E.A. (2012). A map of visual space in the primate entorhinal cortex. Nature 491, 761–764. 10.1038/nature11587.

29. Bao, X., Gjorgieva, E., Shanahan, L.K., Howard, J.D., Kahnt, T., and Gottfried, J.A. (2019). Grid-like Neural Representations Support Olfactory Navigation of a Two-Dimensional Odor Space. Neuron 102, 1066–1075 e1065. 10.1016/j.neuron.2019.03.034.

30. Constantinescu, A.O., O’Reilly, J.X., and Behrens, T.E.J. (2016). Organizing conceptual knowledge in humans with a gridlike code. Science 352, 1464–1468. 10.1126/science.aaf0941.

31. Horner, A.J., Bisby, J.A., Zotow, E., Bush, D., and Burgess, N. (2016). Grid-like Processing of Imagined Navigation. Curr Biol 26, 842–847. 10.1016/j.cub.2016.01.042.

32. Park, S.A., Miller, D.S., and Boorman, E.D. (2021). Inferences on a multidimensional social hierarchy use a grid-like code. Nat Neurosci 24, 1292–1301. 10.1038/s41593-021-00916-3.

33. Jeffery, K.J. (2007). Integration of the sensory inputs to place cells: what, where, why, and how? Hippocampus 17, 775–785. 10.1002/hipo.20322.

34. Bellmund, J.L.S., Gardenfors, P., Moser, E.I., and Doeller, C.F. (2018). Navigating cognition: Spatial codes for human thinking. Science 362. 10.1126/science.aat6766.

35. Buzsaki, G., and Moser, E.I. (2013). Memory, navigation and theta rhythm in the hippocampal-entorhinal system. Nat Neurosci 16, 130–138. 10.1038/nn.3304.

36. Dang, S., Wu, Y., Yan, R., and Tang, H. (2021). Why grid cells function as a metric for space. Neural Netw 142, 128–137. 10.1016/j.neunet.2021.04.031.

37. O’Keefe, J., and Krupic, J. (2021). Do hippocampal pyramidal cells respond to nonspatial stimuli? Physiol Rev 101, 1427–1456. 10.1152/physrev.00014.2020.

38. Itskov, P.M., Vinnik, E., Honey, C., Schnupp, J., and Diamond, M.E. (2012). Sound sensitivity of neurons in rat hippocampus during performance of a sound-guided task. J Neurophysiol 107, 1822–1834. 10.1152/jn.00404.2011.

39. Komorowski, R.W., Manns, J.R., and Eichenbaum, H. (2009). Robust conjunctive item-place coding by hippocampal neurons parallels learning what happens where. J Neurosci 29, 9918–9929. 10.1523/JNEUROSCI.1378-09.2009.

40. Moita, M.A., Rosis, S., Zhou, Y., LeDoux, J.E., and Blair, H.T. (2003). Hippocampal place cells acquire location-specific responses to the conditioned stimulus during auditory fear conditioning. Neuron 37, 485–497. 10.1016/s0896-6273(03)00033-3.

41. Eichenbaum, H., Yonelinas, A.P., and Ranganath, C. (2007). The medial temporal lobe and recognition memory. Annu Rev Neurosci 30, 123–152. 10.1146/annurev.neuro.30.051606.094328.

42. Ranganath, C., and Ritchey, M. (2012). Two cortical systems for memory-guided behaviour. Nat Rev Neurosci 13, 713–726. 10.1038/nrn3338.

43. Burwell, R.D. (2000). The parahippocampal region: corticocortical connectivity. Ann N Y Acad Sci 911, 25–42. 10.1111/j.1749-6632.2000.tb06717.x.

44. Agster, K.L., and Burwell, R.D. (2009). Cortical efferents of the perirhinal, postrhinal, and entorhinal cortices of the rat. Hippocampus 19, 1159–1186. 10.1002/hipo.20578.

45. Burwell, R.D., and Amaral, D.G. (1998). Cortical afferents of the perirhinal, postrhinal, and entorhinal cortices of the rat. J Comp Neurol 398, 179–205. 10.1002/(sici)1096-9861(19980824)398:2<179::aid-cne3>3.0.co;2-y.

46. Todd, T.P., Mehlman, M.L., Keene, C.S., DeAngeli, N.E., and Bucci, D.J. (2016). Retrosplenial cortex is required for the retrieval of remote memory for auditory cues. Learn Mem 23, 278–288. 10.1101/lm.041822.116.

47. Suzuki, W.A., and Amaral, D.G. (1994). Perirhinal and parahippocampal cortices of the macaque monkey: cortical afferents. J Comp Neurol 350, 497–533. 10.1002/cne.903500402.

48. Zhang, G.W., Sun, W.J., Zingg, B., Shen, L., He, J., Xiong, Y., Tao, H.W., and Zhang, L.I. (2018). A Non-canonical Reticular-Limbic Central Auditory Pathway via Medial Septum Contributes to Fear Conditioning. Neuron 97, 406–417 e404. 10.1016/j.neuron.2017.12.010.

49. Koganezawa, N., Gisetstad, R., Husby, E., Doan, T.P., and Witter, M.P. (2015). Excitatory Postrhinal Projections to Principal Cells in the Medial Entorhinal Cortex. J Neurosci 35, 15860–15874. 10.1523/JNEUROSCI.0653-15.2015.

50. Czajkowski, R., Sugar, J., Zhang, S.J., Couey, J.J., Ye, J., and Witter, M.P. (2013). Superficially projecting principal neurons in layer V of medial entorhinal cortex in the rat receive excitatory retrosplenial input. J Neurosci 33, 15779–15792. 10.1523/JNEUROSCI.2646-13.2013.

51. Kerr, K.M., Agster, K.L., Furtak, S.C., and Burwell, R.D. (2007). Functional neuroanatomy of the parahippocampal region: the lateral and medial entorhinal areas. Hippocampus 17, 697–708. 10.1002/hipo.20315.

52. Olsen, G.M., Ohara, S., Iijima, T., and Witter, M.P. (2017). Parahippocampal and retrosplenial connections of rat posterior parietal cortex. Hippocampus 27, 335–358. 10.1002/hipo.22701.

53. Dana, H., Chen, T.W., Hu, A., Shields, B.C., Guo, C., Looger, L.L., Kim, D.S., and Svoboda, K. (2014). Thy1-GCaMP6 transgenic mice for neuronal population imaging in vivo. PLoS One 9, e108697. 10.1371/journal.pone.0108697.

54. Gu, Y., Lewallen, S., Kinkhabwala, A.A., Domnisoru, C., Yoon, K., Gauthier, J.L., Fiete, I.R., and Tank, D.W. (2018). A Map-like Micro-Organization of Grid Cells in the Medial Entorhinal Cortex. Cell 175, 736–750 e730. 10.1016/j.cell.2018.08.066.

55. Obenhaus, H.A., Zong, W., Jacobsen, R.I., Rose, T., Donato, F., Chen, L., Cheng, H., Bonhoeffer, T., Moser, M.B., and Moser, E.I. (2022). Functional network topography of the medial entorhinal cortex. Proc Natl Acad Sci U S A 119. 10.1073/pnas.2121655119.

56. Hardcastle, K., Maheswaranathan, N., Ganguli, S., and Giocomo, L.M. (2017). A Multiplexed, Heterogeneous, and Adaptive Code for Navigation in Medial Entorhinal Cortex. Neuron 94, 375–387 e377. 10.1016/j.neuron.2017.03.025.

57. Fyhn, M., Hafting, T., Treves, A., Moser, M.B., and Moser, E.I. (2007). Hippocampal remapping and grid realignment in entorhinal cortex. Nature 446, 190–194. 10.1038/nature05601.

58. Radvansky, B.A., Oh, J.Y., Climer, J.R., and Dombeck, D.A. (2021). Behavior determines the hippocampal spatial mapping of a multisensory environment. Cell Rep 36, 109444. 10.1016/j.celrep.2021.109444.

59. Geva-Sagiv, M., Romani, S., Las, L., and Ulanovsky, N. (2016). Hippocampal global remapping for different sensory modalities in flying bats. Nat Neurosci 19, 952–958. 10.1038/nn.4310.

60. Bowler, J.C., and Losonczy, A. (2023). Direct cortical inputs to hippocampal area CA1 transmit complementary signals for goal-directed navigation. Neuron. 10.1016/j.neuron.2023.09.013.

61. Davoudi, H.I., J. B.; Climer, J. R.; Dombeck, D. A. (2023). Entorhinal-hippocampal spatial representations during multisensory navigation. held in Washington DC, (Society for Neuroscience).

62. Libby, L.A., Ekstrom, A.D., Ragland, J.D., and Ranganath, C. (2012). Differential connectivity of perirhinal and parahippocampal cortices within human hippocampal subregions revealed by high-resolution functional imaging. J Neurosci 32, 6550–6560. 10.1523/JNEUROSCI.3711-11.2012.

63. Kropff, E., Carmichael, J.E., Moser, E.I., and Moser, M.B. (2021). Frequency of theta rhythm is controlled by acceleration, but not speed, in running rats. Neuron 109, 1029–1039 e1028. 10.1016/j.neuron.2021.01.017.

64. Sheintuch, L., Rubin, A., Brande-Eilat, N., Geva, N., Sadeh, N., Pinchasof, O., and Ziv, Y. (2017). Tracking the Same Neurons across Multiple Days in Ca(2+) Imaging Data. Cell Rep 21, 1102–1115. 10.1016/j.celrep.2017.10.013.

65. Domnisoru, C., Kinkhabwala, A.A., and Tank, D.W. (2013). Membrane potential dynamics of grid cells. Nature 495, 199–204. 10.1038/nature11973.

66. Cholvin, T., Hainmueller, T., and Bartos, M. (2021). The hippocampus converts dynamic entorhinal inputs into stable spatial maps. Neuron 109, 3135–3148 e3137. 10.1016/j.neuron.2021.09.019.

